# Top-Down Proteomics Identifies Plasma Proteoform Signatures of Liver Cirrhosis Progression

**DOI:** 10.1101/2024.06.19.599662

**Authors:** Eleonora Forte, Jes M. Sanders, Indira Pla, Vijaya Lakshmi Kanchustambham, Michael A. R. Hollas, Che-Fan Huang, Aniel Sanchez, Katrina N. Peterson, Rafael D. Melani, Alexander Huang, Praneet Polineni, Julianna M. Doll, Zachary Dietch, Neil L. Kelleher, Daniela P. Ladner

**Affiliations:** Proteomics Center of Excellence, Northwestern University, Evanston, IL, 60208, USA; Northwestern University Transplant Outcomes Research Collaborative (NUTORC), Comprehensive Transplant Center, Feinberg School of Medicine, Northwestern University, Chicago, IL, 60611, USA; Department of Chemistry, Northwestern University, Evanston, IL, 60208, USA; Department of Biochemistry and Molecular Genetics, Northwestern University Feinberg School of Medicine, Chicago, IL, 60611, USA

## Abstract

Cirrhosis, advanced liver disease, affects 2-5 million Americans. While most patients have compensated cirrhosis and may be fairly asymptomatic, many decompensate and experience life-threatening complications such as gastrointestinal bleeding, confusion (hepatic encephalopathy), and ascites, reducing life expectancy from 12 to less than 2 years. Among patients with compensated cirrhosis, identifying patients at high risk of decompensation is critical to optimize care and reduce morbidity and mortality. Therefore, it is important to preferentially direct them towards specialty care which cannot be provided to all patients with cirrhosis. We used discovery Top-down Proteomics (TDP) to identify differentially expressed proteoforms (DEPs) in the plasma of patients with progressive stages of liver cirrhosis with the ultimate goal to identify candidate biomarkers of disease progression. In this pilot study, we identified 209 DEPs across three stages of cirrhosis (compensated, compensated with portal hypertension, and decompensated), of which 115 derived from proteins enriched in the liver at a transcriptional level and discriminated the three stages of cirrhosis. Enrichment analyses demonstrated DEPs are involved in several metabolic and immunological processes known to be impacted by cirrhosis progression. We have preliminarily defined the plasma proteoform signatures of cirrhosis patients, setting the stage for ongoing discovery and validation of biomarkers for early diagnosis, risk stratification, and disease monitoring.

**Graphical Abstract:** 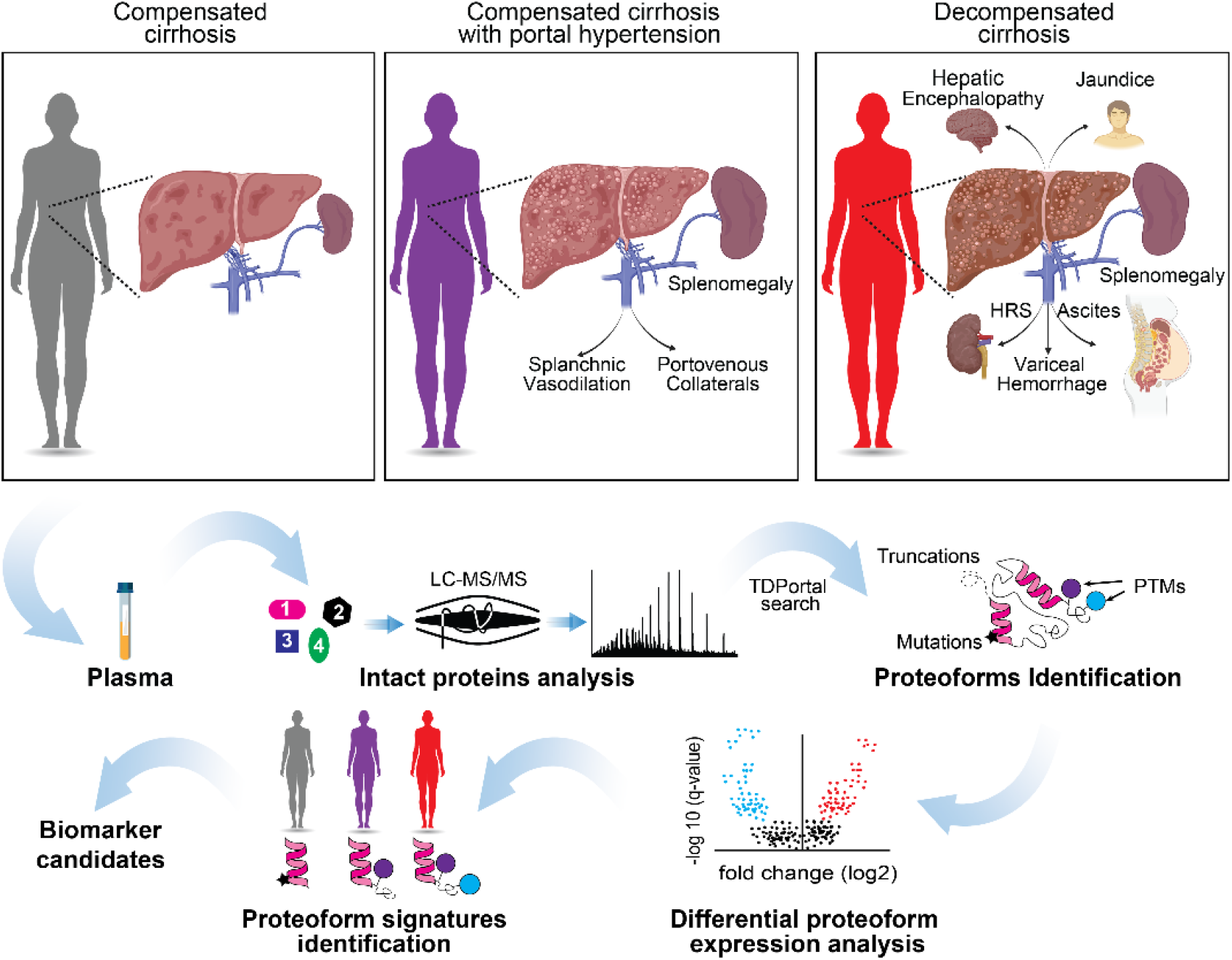

**Highlights:**

- Performed a pilot top-down LC-MS/MS analysis to identify proteoforms (PFRs) in the plasma of patients with 3 progressive stages of liver cirrhosis.
- Identified 2867 proteoforms (PFRs) and 209 differentially regulated proteoforms (DRPs) in the different stages of the disease.
- Identified DRP profiles able to potentially distinguish early from late stages of the disease, including 115 liver-derived DRPs.
- Fibrinogen alpha chain, haptoglobin, and Apo A-I are the proteins with the highest number of DRPs and represent potential candidate biomarkers of liver cirrhosis progression.

## Introduction

The prevalence of cirrhosis and end-stage liver disease is increasing and pose a significant public health burden, in terms of hospitalizations, cost and mortality ^1–5^. In fact, mortality of cirrhosis is comparable to diabetes and pneumonia and surpasses the mortality associated with most cancers ^6, 7^. Cirrhosis is the result of liver injury most frequently caused by alcohol use disorder, metabolic dysfunction-associated liver disease [MASLD; previously, non-alcoholic fatty liver disease (NAFLD)], and hepatitis C (HCV), along with other less frequent etiologies (e.g., biliary, hepatitis B). Most patients with cirrhosis have compensated chronic disease, which carries a life expectancy of ∼12 years ^8^. However, as cirrhosis progresses, intrahepatic structural changes lead to diversion of blood flow to the heart through alternate small blood vessels, rather than the liver. This process of porto-venous collateral formation and splanchnic vasodilation is called portal hypertension (pHTN) ^9–11^. As liver disease progresses, pHTN can turn into decompensated cirrhosis, considered as the occurrence of confusion (hepatic encephalopathy), ascites, and gastrointestinal bleeding. Decompensated cirrhosis is associated with significant morbidity, high rates of hospitalizations, and decreased life expectancy of 1.8 years. On average one in ten patients with cirrhosis experience a decompensating event every year ^8^. With the increasing prevalence of metabolic syndrome and alcohol use disorder, the two major causes of cirrhosis, cirrhosis-related morbidity and mortality are expected to rise ^5, 12^.

Early diagnosis, optimal disease management, and identification of patients who are at high risk of decompensation is critical to provide patients with targeted and timely interventions, mitigate disease progression, and decrease morbidity and mortality ^13, 14^. While only experimental drugs exist to mitigate disease progression ^15, 16^, timely specialty care can optimize the disease course. However, with 2-5M adults affected by cirrhosis, and limited resources including specialists, identifying those at highest risk for disease progression is paramount for optimal triage and outcomes. Presently, such diagnostic tools are not widely available. Hepatic vein pressure gradients (HVPG) have been shown to be a marker of decompensation ^17^, but the procedure is invasive and may not be available at all centers. Clinical risk scores may hold some prognostic value, but the majority were initially established to predict mortality and not disease progression ^18, 19^. To this end, non-invasive and reliable diagnostic tools to identify and predict identify those at highest risk are required. Top-down proteomics (TDP) can be used to develop predictive markers of disease progression by leveraging modified proteins that drive mechanisms of disease. TDP can characterize proteins present in biological systems by determining the molecular composition of all forms of a given protein, so called “proteoforms.” Proteoforms (PFRs) arise due to genetic variation in protein coding regions, alternative splicing, and/or post-translational modifications (PTMs) ^20^. It is estimated that the human proteome contains >50 million unique PFRs compared to ∼20,300 gene-encoded proteins ^21^. As a result, by delivering information at the proteoform level, TDP captures the complexity of a biological system better than that of traditional peptide-based proteomics approaches, which only provide data at the protein level. We have recently demonstrated the importance of PFRs as biomarkers in the context of liver transplantation with the identification of PFR signatures of hematopoietic stem cells able to distinguish acute rejection from normal graft function ^22^. PFRs specific for cirrhosis, especially those specific to different stages of cirrhosis, have not been described, and their identification could greatly improve the diagnosis and treatment of those at highest risk for progression.

In this study, we applied a TDP approach to identify PFRs present in the plasma of patients with three different stages of cirrhosis [compensated (Stage I)/ compensated + pHTN (Stage II) / decompensated (Stage III)]. We found that patients with different stages of cirrhosis have unique blood proteoform profiles that impact important pathophysiological pathways and cellular processes. Our findings create a framework for applying proteoform-based analysis to biomarker discovery in the cirrhosis population and can provide mechanistic insight to the disrupted processes as the disease progresses.

### Experimental Procedures

#### Human Subject Selection and Plasma Collection

Peripheral blood was collected from 30 subjects with cirrhosis as part of clinical cirrhosis care at Northwestern Medicine who had compensated cirrhosis (Stage I), compensated cirrhosis + pHTN (Stage II), and decompensated cirrhosis (Stage III). Plasma was then isolated and stored at -80°C until protein extraction. Samples from adults (>18 years) with 1) compensated cirrhosis, 2) compensated cirrhosis + pHTN, and 3) decompensated cirrhosis were used. Diagnosis of cirrhosis and stage of cirrhosis was determined by careful medical record review (JMS, PP, DPL), including Fibroscan®, magnetic resonance elastography, or biopsy. Decompensation was defined as the presence of hepatic encephalopathy (HE), ascites, hepatorenal syndrome, spontaneous bacterial peritonitis, and history of variceal gastrointestinal bleeding ^23, 24^. pHTN was diagnosed either by imaging, presence of thrombocytopenia (platelets <100), or evidence of esophageal/gastric varices or portal hypertensive gastropathy on surveillance/diagnostic endoscopy ^24, 25^. Demographic, clinical characteristics, and laboratory values (complete blood count (CBC), liver enzymes, basic chemistries, and INR) were collected for each subject at the time of lab draw. (Figure S1). The Model for End Stage Liver Disease Sodium (MELD-Na) was calculated for each subject.

#### Study approval

This study was conducted in compliance with the NIH guidelines for studies involving human subjects. The Northwestern Institutional Review Board (IRB) issued a Waiver of Consent and approved this study under STU00216399.

#### Plasma Protein Extraction and Fractionation

Protein concentration in the plasma was determined by a bicinchoninic acid (BCA) protein assay kit (Thermo Fisher Scientific) following the manufacturer instructions. Passively Eluting Proteins from Polyacrylamide gels as Intact species for MS (PEPPI-MS) was used to fractionate proteins with a molecular weight lower than 30kDa and prepare them for the mass spectrometry analysis ^26^. Briefly, 500 µg of plasma proteins were resuspended in 4X NuPAGE loading buffer (Thermo Fisher Scientific) with 50 mM DTT and boiled at 95°C for 10 minutes. Proteins were fractionated on a NuPAGE 4-12% Bis-Tris gel (Thermo Fisher Scientific) for 10 min at 70 V and 15 minutes at 150 V. The gel was briefly rinsed with deionized water and gel sections with proteins with a molecular weight below 30kDa were excised from the gel and transferred to a Protein LoBind tube (Eppendorf) containing 100 mM ammonium bicarbonate, pH 9 with 0.1% SDS. Proteins were extracted from the gel by crushing it with a pestle and shaking the slurry for 20 minutes at 1400 rpm. Gel debris were removed with 0.45 µm centrifugal filters. The protein fractions were methanol/chloroform/water precipitated by using a slightly modified Wessel and Flügge’s method ^26, 27^. Briefly, 2 volumes of methanol and 0.5 volume of chloroform were added to 1 volume of the protein sample and the mixture vortexed. 1 volume of water was then added to the solution, and the mixture was vortexed and centrifuged at 20,000 rpm for 20 minutes at 4°C. Without disturbing the interface, most of the upper layer was withdrawn and discarded. Next, 1 mL of methanol was added to the lower phase and the mixture was vortexed and centrifuged at 20,000 rpm for 30 minutes. The supernatant was removed. The precipitated protein pellet was air-dried, resuspended in 30 μL of LC-MS buffer A (5% acetonitrile, 94.8% water, and 0.2% formic acid), and subjected to LC-MS/MS.

#### LC-MS/MS Analysis

Proteoforms separation was conducted on a Dionex Ultimate 3000 (Thermo Fisher Scientific) by using a trap/column system consisting of: 1) a trap (150 μm I.D. x 2.5 cm length) packed in-house with ReproSil C4, 3 μm particles (Dr. Maisch), 2) a FlowChip column (25 cm in length x 100 µm i.d. obtained from New Objective). For the separation the flow rate was set at 1 μL/min and the following gradient was used: 5% B (5% water, 94.8% acetonitrile, and 0.2% formic acid) from 0 to 10 min., 20% B at 15 min, 55% B at 100 min, 95% B from 103 to 110 min, 5% B at 115 to 120 min.

The LC is in line with the Orbitrap Eclipse (Thermo Fisher Scientific) mass spectrometer operating in “protein mode” with 2 mTorr of N_2_ pressure in the ion routing multipole (IRM). Transfer capillary temperature was set at 320°C, ion funnel RF was set at 50%, and a 15 V of source CID was applied. MS^1^ spectra were acquired at 120,000 of resolving power (at *m/z* 200), AGC target value of 1000%, 100 ms of maximum injection time, and 1 μscan. Data-dependent top-N-2 sec MS^2^ method used 32 NCE for HCD to generate fragmentation spectra acquired at 60,000 resolving power (at *m/z* 200), with target AGC values of 2000%, 600 ms maximum injection time, and 1 μscan. Precursors were quadrupole isolated using a 3 *m/z* isolation window, dynamic exclusion of 60 s duration, and threshold of 1×10^4^ intensity. Each sample was acquired in triplicate for a total of 90 runs.

### Experimental Design and Statistical Rationale

#### Data Analysis for LC-MS/MS

The raw data files were processed with the publicly available workflow on TDPortal (https://portal.nrtdp.northwestern.edu, Code Set 4.0.0) that performs mass inference, searches a database of human proteoforms derived from Swiss-Prot (June 2020) with curated histones, and estimates a conservative, context-dependent 1% false discovery rate (FDR) at the protein, isoform, and proteoform levels ^28^.

#### Statistics and Functional enrichment analysis

Data derived from randomized biological & technical replicates were used to assess variability associated with sample processing. From the aggregated data sets a batch effect was observed and corrected with batch standardization (z-score) and statistical analysis was conducted using SAS (Cary, NC) with custom scripts. The appropriate hierarchical linear statistical model was applied for quantitation using SAS PROC MIXED (SAS Institute). Normalized and batch corrected ion intensities from the data files were run through a nested ANOVA process to measure both effect size and significance in differential proteoform expression between the 3 patients’ groups. Specific proteoform signatures were then correlated with decompensated outcomes. P-values obtained from the mixed model linear regression were adjusted (Benjamini & Hochberg (BH) algorithm) to control the false discovery rate (FDR) induced by multiple comparison tests. PFRs with adjusted p value < 0.05 and fold changes higher than 1.5 or less than 0.66, were considered differential expressed proteoforms (DEP). A Partitioning Around Medoids (PAM) ^29^ unsupervised algorithm (‘cluster::Pam’Rfunction) was applied to group PFRs based on their expression changes between different disease stages. The number of clusters (k) was manually set to three, as it was the number of clusters that best showed changes in disease progression. To visualize clusters based on differential expressions and changes in specific PFR signatures we created heatmaps using the ‘ComplexHeatmap’ R package. Clinical biomarkers included in heatmap from Figure 2 were previously standardized across samples using z-score normalization. To know whether the PFR modifications were significantly different between groups of DEPfr, we applied Fisher exact tests followed by an adjustment of p-values (BH). Proteins having at least one deregulated PFR were selected to perform Gene Ontology enrichment analysis based on Biological Process, and the analysis was carried out using ‘clusterProfiles’ R package.

## Results

### 1.1 Discovery Top-Down Proteomics (TDP) analysis of plasma characterizes blood proteoform profiles in cirrhosis patients

The cohort included 30 subjects—9 (30%) with compensated cirrhosis (Stage I), 10 (33.3%) with compensated cirrhosis + pHTN (Stage II), and 11 (36.7%) with decompensated cirrhosis (Stage III) (Figure 1A). Demographics and clinical data are shown in Table 1. The mean age of patients was 59.7 years (Stage I: 63.7, Stage II: 55.5, Stage III: 61.9). 40% (n=12) of patients were female (Stage I: 55.6%, Stage II 40.0%, Stage III: 27.3%), and 56.7% (n=17) were White. The main etiology of cirrhosis was alcohol use disorder in 60% (n=18). The mean Model for End-stage Liver Disease – Sodium (MELD-Na) was 11.5 (Stage I: 9.9, Stage II: 10.0, and Stage III: 14.1). Detailed laboratory findings are shown in Figure S1. No clinically significant differences were observed.

**Figure 1.**
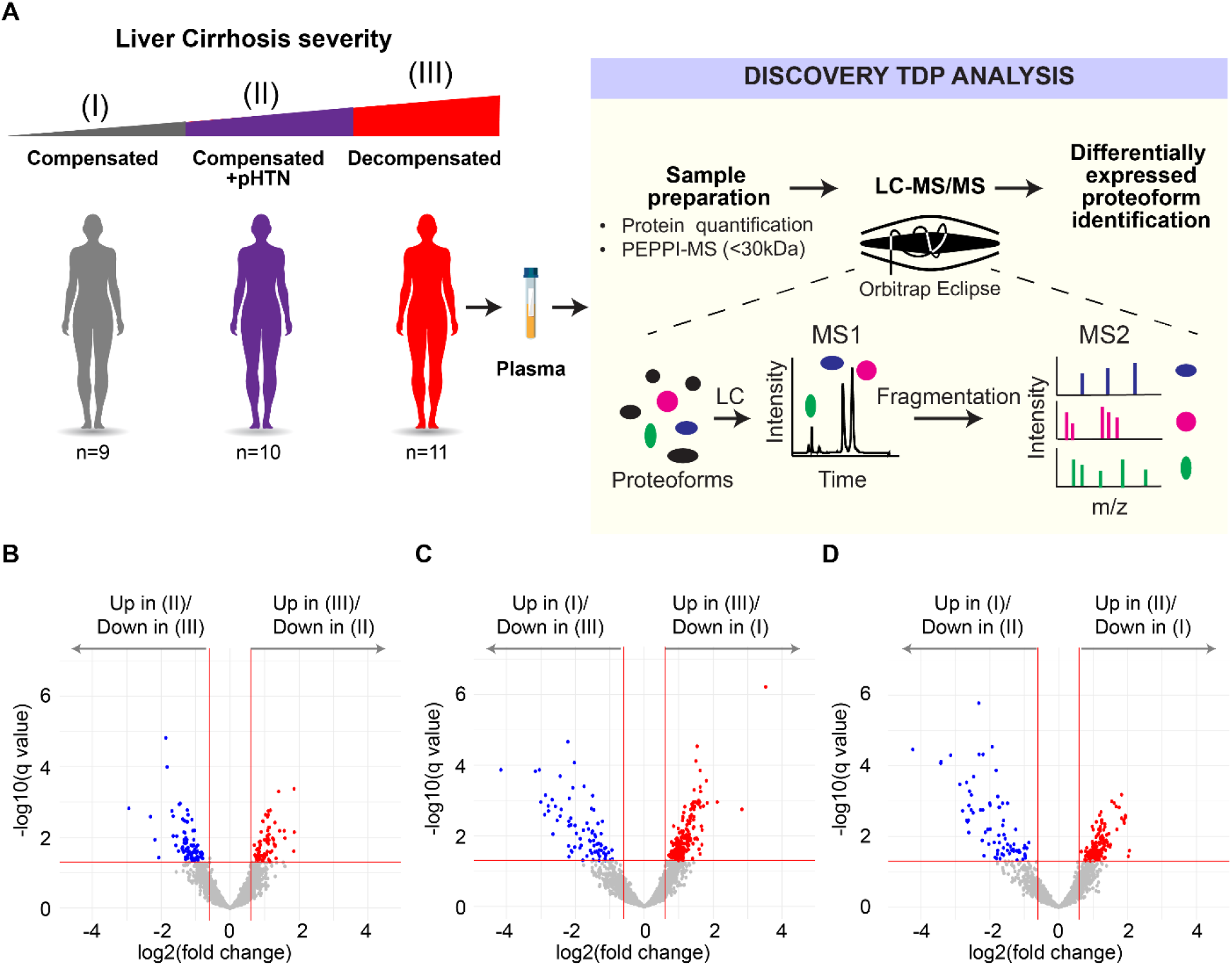
Workflow used to identify differentially expressed proteoforms (DEPs) in patients with cirrhosis. **A)** Plasma was extracted from cirrhosis patients with three stages of disease— compensated (I), compensated with portal hypertension (II), and decompensated cirrhosis (III). 30kDa fractions of plasma were then subjected to Top-Down Proteomics (TDP) to identify DEPs across disease stages. **B-D)** Volcano plots showing differentially expressed proteoforms (DEPs) in the plasma from patients with **B)** decompensated cirrhosis (III) vs compensated cirrhosis with portal hypertension (II), **C)** decompensated cirrhosis (III) vs compensated cirrhosis (I), and **D)** compensated cirrhosis with portal hypertension (II) vs compensated cirrhosis (I). A false discovery rate (FDR) threshold of 5% for differential expression was used.

**Table 1.** Characteristics of the patient cohort. pHTN= portal hypertension, AIH= autoimmune hepatitis, PBC= primary biliary cholangitis, ETOH= alcohol-associated liver disease, HBV= hepatitis B virus, HCV= hepatitis C virus, MASH= metabolic dysfunction associated steatohepatitis, BMI= body mass index, MELD-Na= Model for End-Stage Liver Disease with Sodium.

|  | Total (n=30) | Compensated (n=9) | Compensated w/ pHTN (n=10) | Decompensated (n=11) | p-value |
| --- | --- | --- | --- | --- | --- |
| **Age, mean (SD)** | 59.73 (12.34) | 63.67 (11.18) | 55.51 (10.54) | 61.89 (14.18) | 0.314 |
| **Female, n (%)** | 12 (40) | 5 (55.6) | 4 (40.0) | 3 (27.3) | 0.612 |
| **Race/Ethnicity, n (%)** |  |  |  |  | 0.206 |
| **Asian** | 0 (0.0) | 0 (0.0) | 0 (0.0) | 0 (0.0) |  |
| **Black or African American** | 3 (10) | 2 (22.2) | 1 (10.0) | 0 (0.0) |  |
| **Hispanic or Latino** | 10 (33.33) | 4 (44.4) | 3 (30.0) | 3 (27.3) |  |
| **other** | 2 (6.67) | 0 (0.0) | 0 (0.0) | 2 (18.2) |  |
| **White** | 17 (56.67) | 3 (33.3) | 6 (60.0) | 8 (72.7) |  |
| **Etiology, n (%)** |  |  |  |  | <0.01 |
| **AIH** | 3 (10) | 0 (0.0) | 2 (20.0) | 1 (9.1) |  |
| **PBC** | 1 (3.33) | 0 (0.0) | 0 (0.0) | 1 (9.1) |  |
| **ETOH** | 18 (60.0) | 5 (55.6) | 5 (50.0) | 8 (72.7) |  |
| **HBV** | 1 (3.33) | 1 (11.1) | 0 (0.0) | 0 (0.0) |  |
| **HCV** | 8 (26.67) | 4 (44.4) | 3 (30.0) | 1 (9.1) |  |
| **MASH** | 4 (13.33) | 1 (11.1) | 1 (10.0) | 2 (18.2) |  |
| **BMI, mean (SD)** | 29.14 (5.55) | 31.14 (6.49) | 28.63 (3.21) | 27.99 (6.42) | 0.336 |
| **MELD-Na, mean (median [IQR])** | 11.47 (4.62) | 9.89 (3.18) | 10.00 (3.94) | 14.09 (5.30) | 0.162 |

We first conducted an untargeted quantitative TDP analysis of 0-30 kDa fractions of plasma (Figure 1A). Within the entire cohort 2867 proteoforms from 99 proteins were captured (Table S1). Modifications of proteoforms mainly included C- and N-terminus truncations (Figure S2). Phosphorylation was the most frequent post-translational modification (PTM) (Figure S2). Other common PTMs included monoacetylation, alpha-amino acetylation, S-nitrosyl-L-cysteine, n-pyruvic acid 2-iminyl-L-valine, monomethylation, monohydroxylation, L-gamma-carboxyglutamic acid, and 2-pyrrolidone-5-carboxylic acid (Figure S2).

### 1.2 Proteoforms are differentially expressed in stages of cirrhosis and are involved in biologically and clinically relevant processes

We then separated samples according to their diagnostic stage (I-III) (Figure 1B-1D) and identified a total of 663 differentially expressed proteoforms (DEPs) from 48 proteins. Like the ∼3000 total proteoforms, modifications of DEPs mainly included truncations, phosphorylation, and acetylation (Figure S2). Pairwise comparisons between the three stages of cirrhosis showed: 1) 74 upregulated and 83 downregulated DEPs comparing Stage III to Stage II (Figure 1B), 216 upregulated and 79 downregulated DEPs comparing Stage III to Stage I (Figure 1C), and 136 upregulated and 75 downregulated DEPs comparing Stage II to Stage I (Figure 1D). In our differential expression analysis, we identified several proteoforms from hemoglobin subunits. As hemoglobin presence could be due to sample handling and preparation, hemoglobin proteoforms were omitted from further analyses. In the pairwise comparisons shown in Figures S3, S4, and S5, we found that some PFRs from a single protein were upregulated while other PFRs from the same protein were downregulated. For example, we identified a total of 14 DEPs for fibrinogen alpha chain, of which 5 were downregulated and 9 were upregulated in Stage III vs Stage II (Figure S3), 11 DEPs for Apolipoprotein A-I, of which 2 were downregulated and 9 were upregulated in Stage III vs Stage I (Figure S4), and 2 DEPs for ceruloplasmin, of which one was upregulated and one was downregulated in Stage II vs Stage I (Figure S5). We next analyzed the DEP modifications found in the three pairwise comparisons (Figure S6A). DEP modifications in the comparison between Stages III and I included mostly truncations (83%), followed by monohydroxylation (7%), alpha-amino acetylation (6%), and phosphorylation (2%). Both L-gamma-carboxyglutamic acid and 2-pyrrolidone-5 carboxylic acid residues occurred in 1% of the modifications. In the comparison between Stages II and I, the top three modifications were again truncation (75%), monohydroxylation (11%), and alpha-amino acetylation (9%). Phosphorylated, monomethylated, L-gamma-carboxyglutamic acid and 2-pyrrolidone-5 carboxylic acid residues each occurred in 1% of the modifications. Finally, in the comparison between Stages III vs II, we found 63% of PTMs were truncations followed by phosphorylation (11%), alpha-amino acetylation (11%), monohydroxylation (9%), carboxylation of glutamic acid residues (4%), monoacetylation (1%), and pyrrolidone-5 carboxylic acid residues (1%). We then compared the number of each modification in the three sets of DEPs (sets: Stage III vs I, II vs I, III vs II) and found that phosphorylation was the only PTM to be statistically significantly different between the three sets (Figure 6B).

DEPs were then clustered based on their regulation patterns in each stage (Figure 2A-2B). Laboratory test values from clinical evaluation were also included in the clustering to find possible associations between DEPs and clinical markers of disease severity. We also performed gene ontology (GO) based on biological processes (BP) enrichment analyses to correlate PFR profiles with affected biological processes. Cluster 1 (C1) included DEPs that were significantly upregulated in decompensated disease (Stage III). This cluster includes 57 DEPs from 23 proteins. The proteins with the highest number of DEPs in C1 were fibrinogen alpha chain with 15 DEPs, apolipoprotein A-I with 10 DEPs, and insulin-like growth factor-binding protein 4 with 5 DEPs. Pathway enrichment analysis indicated that C1 DEPs derived from proteins involved in several biological processes, including the humoral immune response, apoptosis regulation, complement activation, phagocytosis, and negative regulation of peptidase activity (Figure 2C, Table S2). In addition, C1 DEPs clustered with several clinical markers, including alkaline phosphatase (ALP), which was significantly higher in Stage III (Figure S1), total bilirubin (TBILI), which was significantly increased between Stage II and III, and MELD-Na, which was significantly and appropriately higher in Stage III compared to Stage I (MELD-Na is a marker of disease severity). Other clinical markers included in this cluster were International Normalized Ratio (INR), prothrombin (PT), aspartate aminotransferase (AST), and creatinine (CR), although their increase in Stage III was not statistically significant.

**Figure 2.**
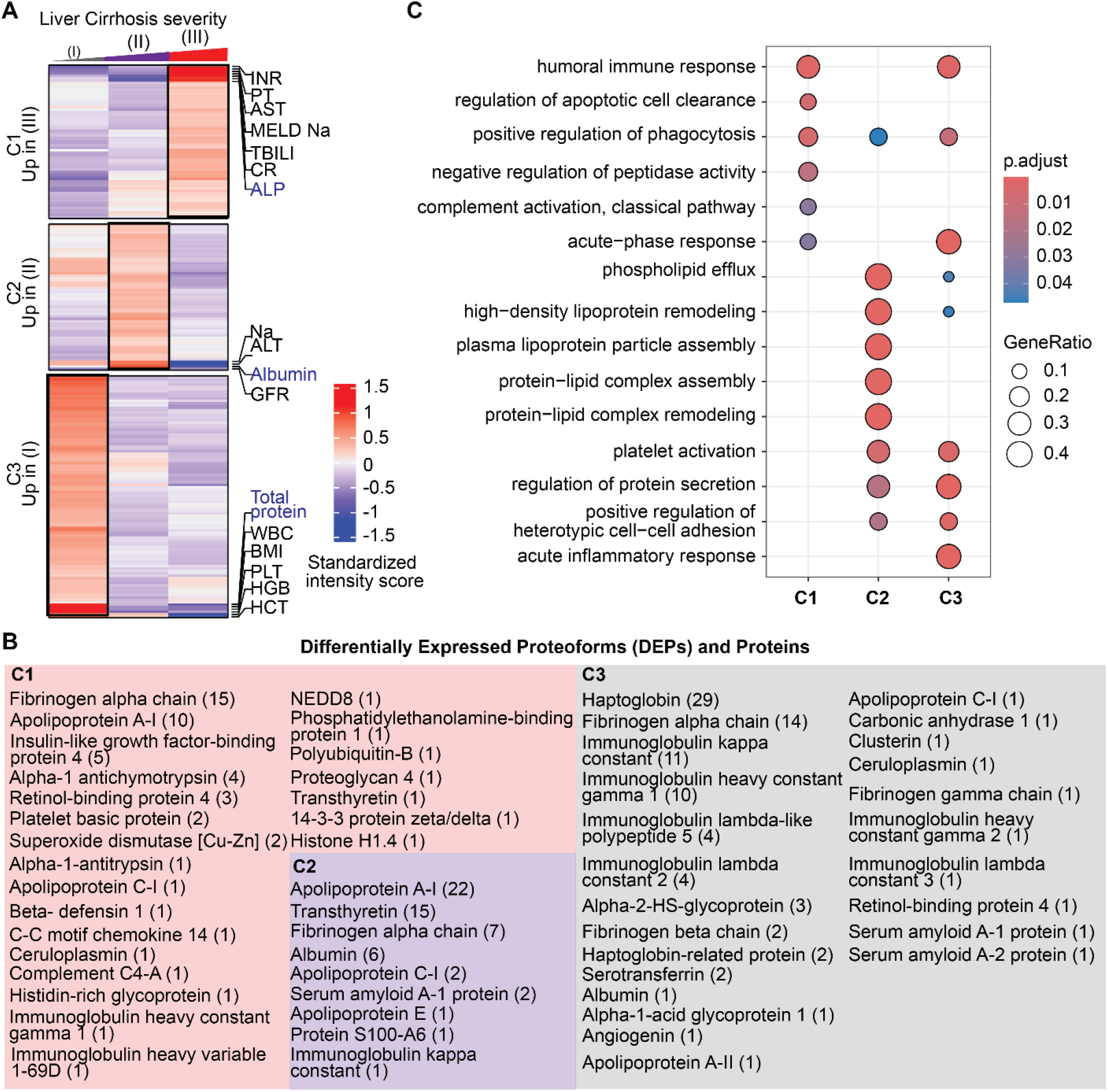
Proteoforms found to be differentially expressed across stages of cirrhosis. **A)** Cluster distribution of differentially expressed proteoforms (DEPs) and currently used clinical markers of disease severity. Clinical markers highlighted in blue are significantly different between stages. Cluster 1 (C1) included DEPs upregulated in Stage III disease, C2 included DEPs upregulated in stage II disease, and C3 included DEPs upregulated in Stage I disease. **B)** Proteins from which DEPs derived are shown within each cluster. The number of proteoforms of a given protein is shown in parentheses beside the protein name. For example, in C1 the fibrinogen alpha chain protein had 15 DEPs identified. **C)** Biological process enrichment analyses on proteins identified in each cluster. The level of significance is represented by the color of the bar with scale while the gene ratio is indicated by the diameter of the circle.

Cluster 2 (C2) included 57 DEPs upregulated in Stage II. The PFRs derived from 9 proteins involved in phospholipid efflux and processes regulated by apolipoproteins (Figure 2C, Table S2). The proteins with the highest number of DEPs were apolipoprotein A-I, transthyretin, and fibrinogen alpha chain with 22, 15, and 7 DEPs, respectively. Clinical markers in this cluster included glomerular filtration rate (GFR), Sodium (Na), alanine aminotransferase (ALT), and albumin. Interestingly, 6 out of 7 albumin PFRs detected in the TDP discovery analysis were found in this cluster together with albumin protein, which was analyzed by routine clinical laboratory testing (Figure S1). Both albumin PFR and protein levels were significantly higher in the plasma of patients with Stage II and lower in those with Stage III. In addition, fold changes obtained for the albumin protein (laboratory results) and albumin PFRs were comparable despite using two distinct approaches, confirming the validity of the TDP discovery analysis.

Cluster 3 (C3) included 95 DEPs upregulated in Stage I. These DEPs derived from 24 proteins associated with the immune system biological processes, such as the acute inflammatory response, acute-phase response, and platelet activation. Other processes enriched in C3 were regulation of protein secretions and cell-cell adhesion (Figure 2C, Table S2). In this cluster, haptoglobin was the protein with the highest number of DEPs (29) followed by fibrinogen alpha chain (16). Interestingly, clinical markers related to the immune system, such as white blood cell count (WBC), platelet count (PLT), and total proteins were grouped in C3. However, only the total protein value changed significantly between Stages I vs II and I vs III (Figure S1).

Overall, clusters containing DEPs upregulated in the late stages of the disease (C1 and C2) were mainly from fibrinogen alpha chain, transthyretin, and apolipoprotein A-I proteins (Figure 2B). However, another important group of DEPs from fibrinogen alpha chain (14 PFRs) were upregulated in Stage I and clustered with haptoglobin in C3. In terms of biological processes, DEPs upregulated in the early stage of the disease were involved in cellular defense mechanisms, while those upregulated in the late stages of cirrhosis were associated with cell death. Percentages of DEP modifications differed in the three clusters as shown in Figure S7A. DEPs in C3 had the highest percentage of truncations (90% versus 77% and 54% of DEPs observed in C1 and C2, respectively). Phosphorylation was higher in C2 (10%) than C1 (3%) and not observed in DEPs in C3. In addition, C2 had a higher percentage of alpha-amino acetylated (C1=6%, C2=13%, C3=6%), monohydroxylated (C1=11%, C2=17%, C3=1%), and L-gamma carboxyglutamic acid (C1=0%, C2= 6%, C3=0%) residues. C2 did not contain any 2-pyrrolidone-5-carboxylic acid modifications, which was instead observed in 1% of C1 and C3 DEPs. Monohydroxylation was only observed in C1 and monomethylation in C3. Finally, the comparison of the number of each modification in the three clusters indicated a significant variation in the types of PFR modifications at the level of phosphorylated, monohydroxylated, alpha-amino acetylated, and L-gamma-carboxyglutamic residues (Figure S7B).

### 1.3 Liver-derived proteoforms may be candidate biomarkers for cirrhosis progression

Heatmaps were generated for the DEPs identified above, but given that cirrhosis is associated with ongoing damage to hepatocytes, we primarily focused on PFRs derived from proteins that are enriched in the liver tissue (https://www.proteinatlas.org) at a transcriptional level as they may be the most sensitive to differences between stages (Figures S3-S5). Heatmaps of these liver-enriched DEPs were created for each stage of cirrhosis, along with their modifications, including truncations and a variety of PTMs (Figure 3). As the majority of DEPs discriminating the different stages of the disease derived from fibrinogen alpha chain, apolipoprotein A-1, and haptoglobin, we analyzed those PFRs in more detail (Figure 3, Figure S8). Fibrinogen alpha chain PFRs were found in all clusters. Specifically, 14 PFRs were detected in C1, 7 in C2 and 15 in C3. The canonical sequence of fibrinogen alpha chain includes 866 amino acids. All the identified PFRs were truncated forms of the protein missing both the N- and C termini. In C1 (PFRs upregulated in Stage III) PFR sequences were generally longer with an average of 126 amino acids. The average length in the other clusters was 58.86 and 58 for C2 and C3, respectively. This suggests the presence of shorter truncations in early stages of the disease. In terms of PTMs, the majority was found in C1. In this cluster we detected 3 PFRs with monohydroxylation and 3 phosphorylated PFRs. In C2, PTMs included one monohydroxylation and one phosphorylation. C3 (PFRs upregulated in Stage I) contained one PFR with monohydroxylation. Of the 32 apolipoprotein A-I PFRs, 22 were in C2 and 10 in C3. The majority were truncated and had 1) no additional PTM, 2) alpha-amino acetylation, or 3) methionine oxidation. A proline 27 to histidine point mutation was also found in 2 PFRs in C2. Haptoglobin PFRs included both truncations of the canonical protein (P00738-1) and of an isoform that lacks amino acids 38-96 (P00738-2). In addition, we found mutations in 10 truncated PFRs of haptoglobin P00738-2, of which eight had an aspartic acid residue replacing asparagine in position 70, and two with lysine at the place of glutamic acid in position 71. In terms of PTMs, we found that four of the haptoglobin PFRs were acetylated at the N-terminus. Although not described in detail, we also identified DEPs deriving from proteins that are not enriched in the liver tissue at a transcriptional level (Figures S3-S5). Heatmaps of non-liver enriched PFRs are displayed in Figure S9.

**Figure 3.**
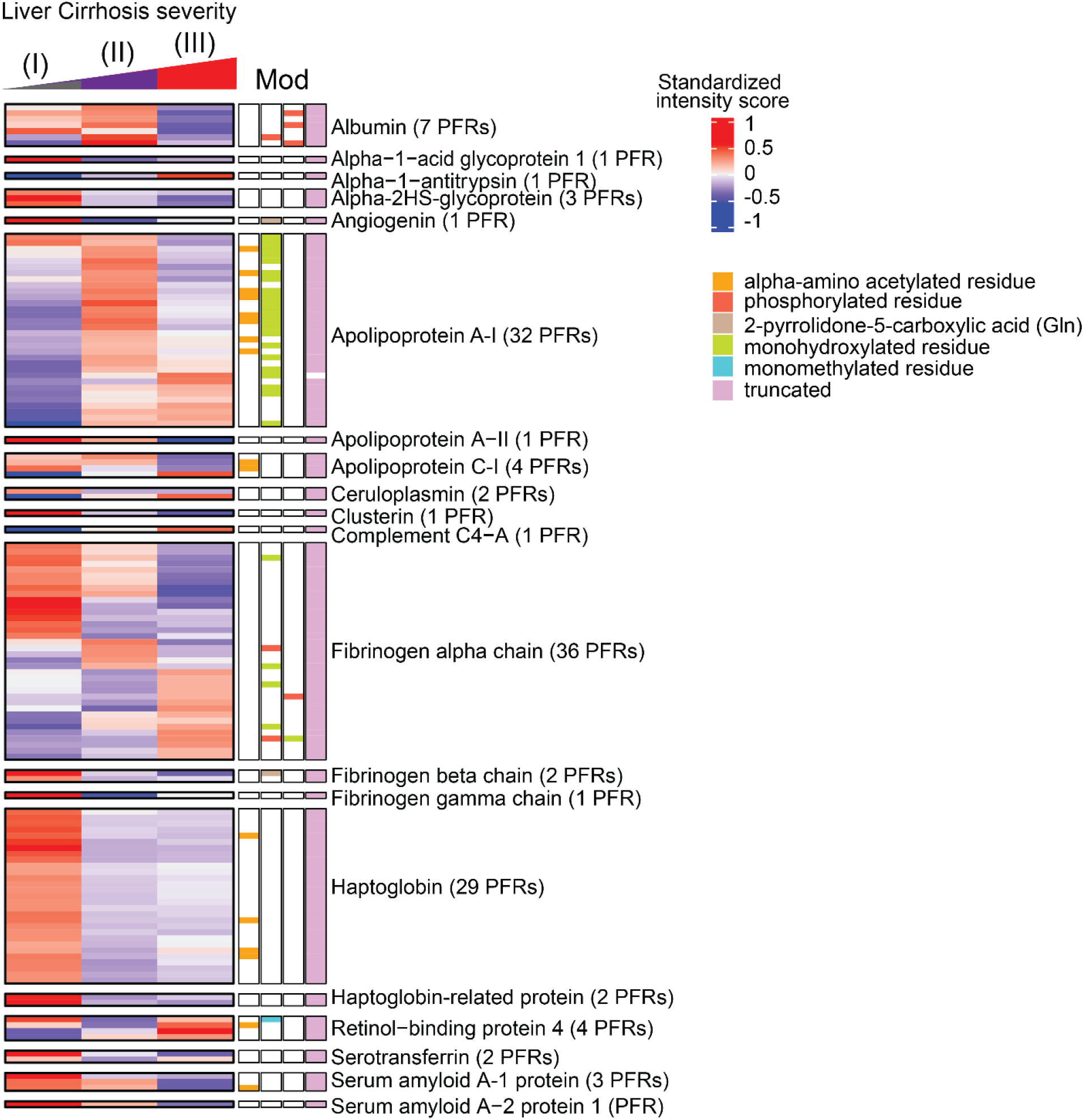
Heatmaps of quantified proteoforms of proteins enriched in the liver at transcriptional level. Associated proteoform modifications are shown and defined by the figure legend. The proteins and number of identified proteoforms derived from each protein are shown to figure right. Abbreviations: PFR (proteoform), MOD (Modification).

To more clearly define PFR signatures that correlated with disease progression, we created profiles of DEPs that could discriminate Stage I (compensated) from Stages II and III of cirrhosis. Comparisons of PFR expression were performed across cirrhosis stages, with compensated disease (Stage I) used as the reference group (Figure 4A). This analysis identified 167 DEPs (from 37 proteins), of which 115 DEPs derived from 20 proteins enriched in the liver at a transcriptional level (Figure 4B). Tables containing detailed information about these 115 DEPs and their modifications are included in Figures S10-12 and Table S1. Apolipoprotein C-I-PFR#16559 was the only DEP found downregulated in Stage III but upregulated in Stages I and II (Figure 4A-4B, and S10, Dark purple square). There were 14 DEPs downregulated in Stage III of the disease compared to Stage I. These included 6 DEPs from fibrinogen alpha chain, 2 from apolipoprotein A-I and serum amyloid A-1 protein, and one from albumin, apolipoprotein A-II, apolipoprotein C-I, and serum amyloid A-2 protein (Figure 4A-4B, Light purple square). Both apolipoprotein A-I PFRs contained oxidized methionine residues at positions 110 and 136 and had N-terminus truncations at two different positions (residues 67 and 35, respectively) (Figure S10). None of the fibrinogen alpha chain PFRs had a PTM, but all contained both C- and N-termini truncations. The average length of truncation was 55.83 amino acids. Three DEPs were specifically downregulated in Stage II disease and derived from serotransferrin, retinol-binding protein 4, and fibrinogen gamma chain (Figure 4A-4B, and S10, Yellow square).

**Figure 4.**
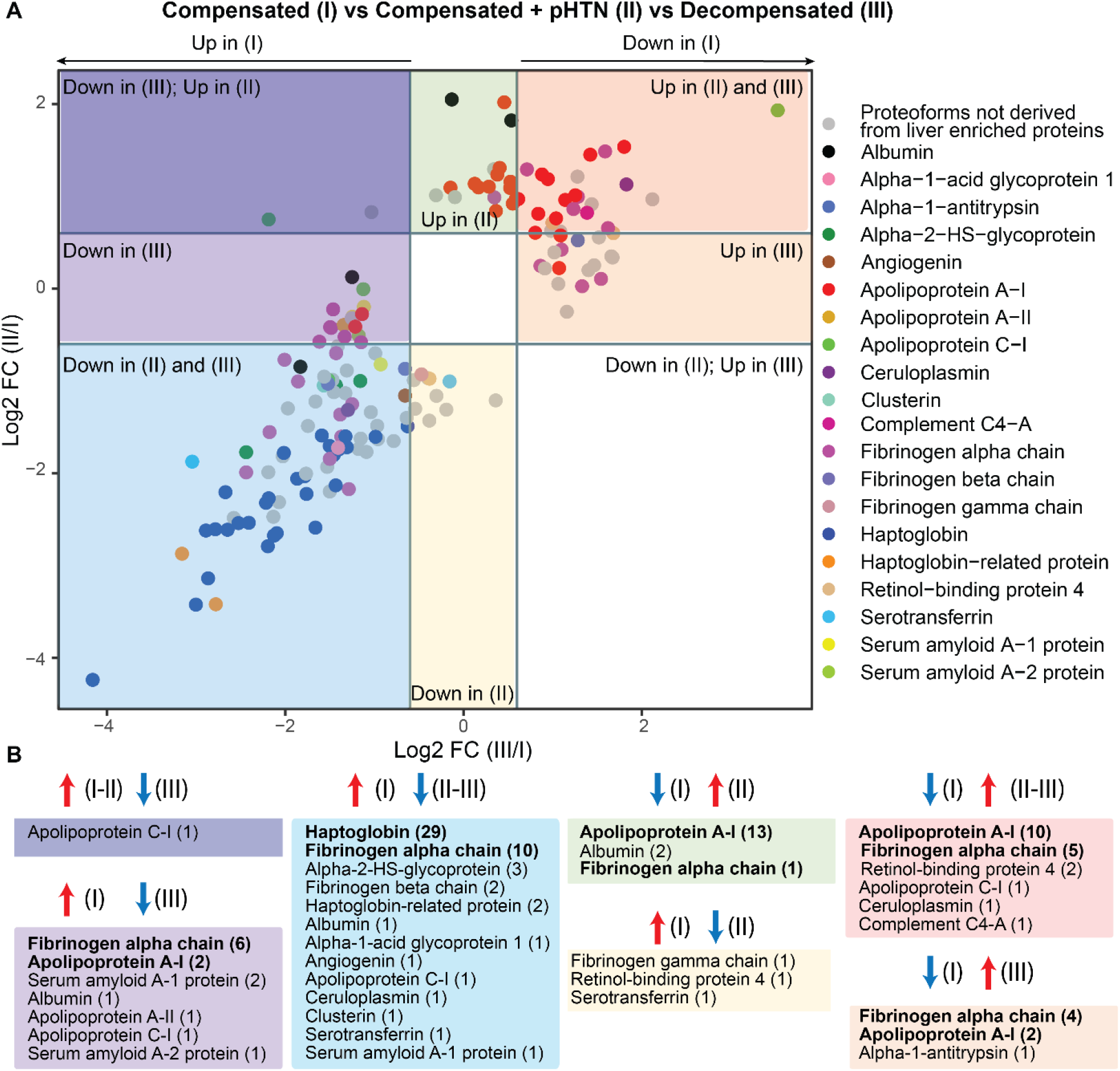
Display of proteoforms stratified by expression in each stage of cirrhosis. **A)** Differentially expressed proteoforms (DEPs) in Stages II and III disease with Stage I disease (compensated) used as the reference. Proteoforms upregulated in Stage II and III (red square), upregulated only in Stage III (orange square), downregulated in Stages II and III (blue square), downregulated in Stage III only (light purple square), downregulated in Stage III but upregulated in Stage II (dark purple square), upregulated only in Stage II (green square), and downregulated only in Stage II (yellow square). Proteoforms deriving from proteins enriched in liver are highlighted and grouped by color according to the figure legend. **B)** Profiles of proteoforms upregulated in Stages I and II and downregulated in III (dark purple), upregulated in I and downregulated in III (light purple), upregulated in I and downregulated in II and II (light blue), downregulated in I and upregulated in II (green), upregulated in I and downregulated in II (yellow), downregulated in I and upregulated in II and III (red), and downregulated in I and upregulated in III (orange). The proteins from which DEPs derive and the relative number of proteoforms (in parenthesis) are shown. Proteins with the highest number of DEPs for each group are shown in bold.

54 DEPs were downregulated in both Stages II and III of the disease compared to Stage I (Figure 4A-4B, and S11, Blue square). Haptoglobin was the protein with the highest number of DEPs (29 PFRs), followed by fibrinogen alpha chain (10 PFRs), alpha-2HS-glycoprotein (3 PFRs), and haptoglobin-related protein (2 PFRs). One DEP each was found for albumin, alpha-1-acid glycoprotein 1, angiogenin, apolipoprotein C-I, ceruloplasmin, clusterin, serotransferrin, and serum amyloid A-1 protein. Haptoglobin PFRs (Figure S11) perfectly overlapped with those characterized in C3 (described above). We again observed both C- and N-termini truncations in all 10 fibrinogen alpha chain PFRs at various positions. The average length of truncation was 58.8 amino acids. One fibrinogen alpha chain PFR was hydroxylated at a proline in position 565 (Figure S11).

Twenty DEPs were upregulated in late Stages (II and III) (Figure 4A-4B, and S12, Red square). These originated from apolipoprotein A-I (10 PFRs), fibrinogen alpha chain (5 PFRs), retinol-binding protein 4 (2 PFRs), apolipoprotein C-I (1 PFR), ceruloplasmin (1 PFR), and complement C4-A (1 PFRs). Sixteen DEPs were exclusively upregulated in Stage II of disease. The upregulated DEPs (Figure 4A-4B, and S12, Green square) were from apolipoprotein A-I (13 PFRs), albumin (2 PFRs), and fibrinogen alpha chain (1 PFR). Finally, 7 DEPs were upregulated in Stage III of the disease and included 4 PFRs derived from fibrinogen alpha chain, 2 from apolipoprotein A-I, and one from alpha-1 antitrypsin (Figure 4A-4B, and S12, Orange square). Collectively, 25 of the 27 apolipoprotein A-I PFRs that discriminated early Stage (I) from later Stage disease (II and III) were upregulated in later disease Stages (Figure 4, Figure S12). These apolipoprotein A-I PFRs mostly contained N-terminus truncations at various locations and oxidation of methionine residues in positions 110 and 136. In addition, a proline to histidine point mutation occurred at position 27 in one PFR that was upregulated only in Stage II disease (Figure S12, green square, PFR#5052090). Notably, all 25 of these PFRs were greater in length compared to those upregulated in Stage I disease (Figure S10). Fibrinogen alpha chain PFRs upregulated in later-stage disease were all truncated at various residues at the N-terminus, but eight of the 10 PFRs had a C-terminal truncation at residue 629. Finally, two fibrinogen alpha chain PFRs were hydroxylated at a proline in position 565 (Figure S12, green and red square), and one was hydroxylated at a serine in position 501 and had a Phospho-proline at position 565 (Figure S12, orange square, PFR#7501917). The average length of truncation of fibrinogen alpha chain PFRs upregulated in later disease stages was higher compared to early-stage disease (Stage III= 137.75, Stages II and III = 73.8, Stage II= 61, Stage I= 55.83-58.8) (Figures 5, S10, S11).

**Figure 5.**
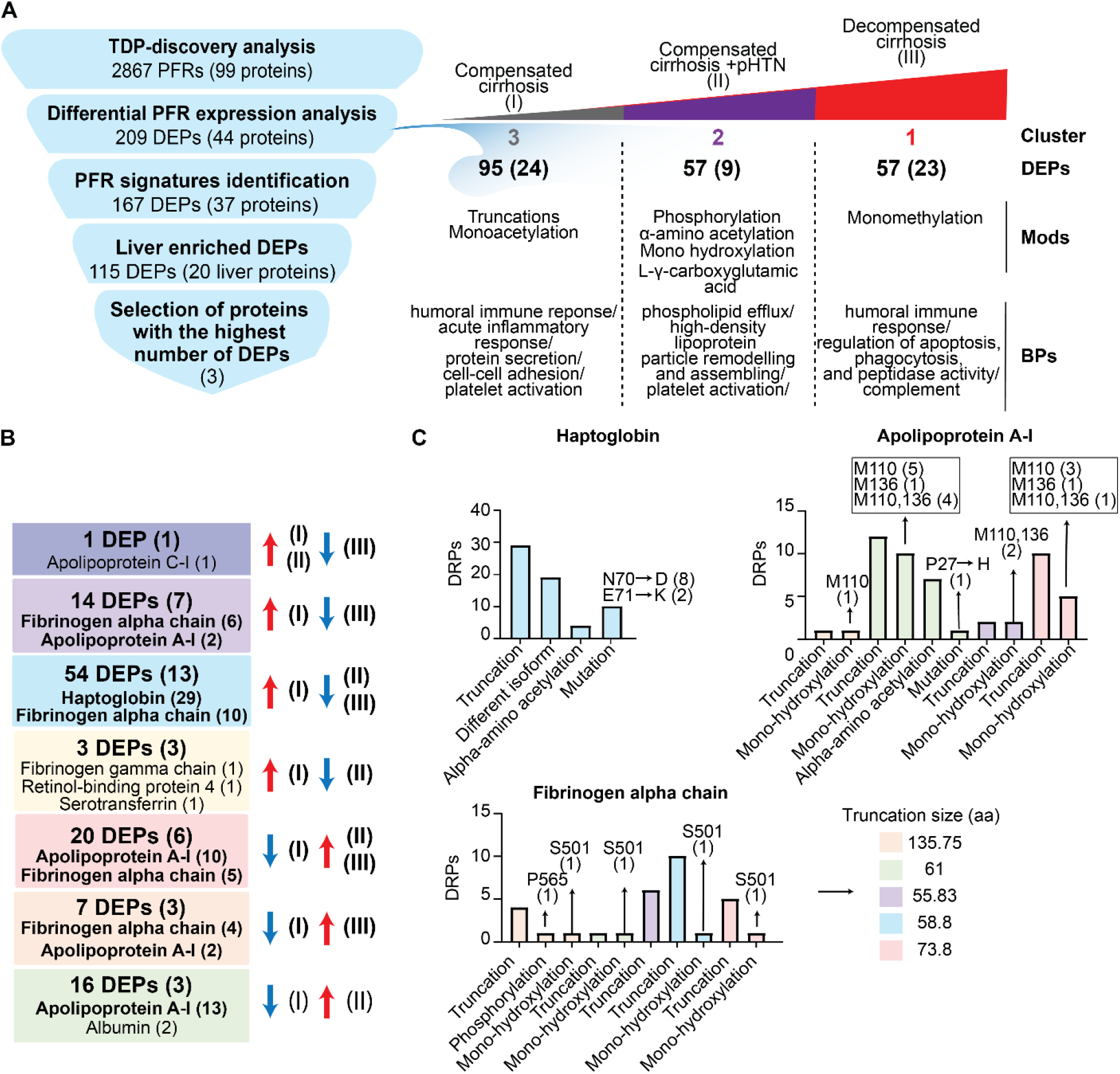
Summary of quantitative TDP analysis of plasma proteoforms in patients with cirrhosis. **A)** Strategy used to identify potential candidate proteoform biomarkers of liver cirrhosis and summary of the differential PFR expression analysis. Differentially expressed proteoforms (DEPs) were classified in clusters based on their upregulation in 1) compensated cirrhosis (Stage I-Cluster 3), 2) compensated cirrhosis with portal hypertension (Stage II-Cluster 2), and 3) decompensated cirrhosis (Stage III-Cluster 1). For each cluster, the number of upregulated DEPs with their relative proteins (in parenthesis), modifications, and biological processes are reported. **B)** Composition of the identified liver derived DEP signatures. Proteins from which the DEPs derive are shown in parenthesis. Proteins with the highest number of DEPs for each group are also reported and shown in bold. DEPs are grouped by color based on their upregulation or downregulation in disease stages as performed in Figure 4. For example, in the red group 20 DEPs deriving from 6 proteins were upregulated in Stages II and III of the disease as compared to Stage I disease. 10 of these DEPs derived from apolipoprotein A-I and 5 derived from fibrinogen alpha chain. **C)** DEPs relative to Haptoglobin, Apolipoprotein A-I, and Fibrinogen alpha chain, which are the 3 plasma proteins with the highest number of detected DEPs, are reported together with their modifications. Colors indicate the signature groups in B and C. Abbreviations: PFR (proteoform), DEP (Differentially expressed proteoform), Mod (Modification) BP (biological process), aa (amino acids).

## Discussion

Using quantitative discovery TDP, we discovered DEPs in three stages of cirrhosis – compensated (Stage I), compensated + pHTN (Stage II) and decompensated (Stage III) (Figure 5). Our pilot discovery analysis identified 2867 PFRs in total, of which 663 were differentially expressed. DEPs clustered into 3 groups based on their upregulation in the various stages of cirrhosis. Modifications to these DEPs differed depending on the stage of cirrhosis, and DEPs were involved in numerous, biologically relevant processes. 167 DEPs discriminated Stage I vs Stages II and III, of which 115 derived from 20 proteins enriched in the liver at a transcriptional level. Proteoforms originating from fibrinogen alpha chain, apolipoprotein A-I, and haptoglobin were the most abundant and demonstrated varying degrees of modification across disease stages.

Apolipoprotein A-I is a lipoprotein produced by both the liver and intestine and is a key component of high-density lipoprotein (HDL). Its primary function is in reverse cholesterol transport, but it is also an important mediator of immunological and inflammatory processes ^30^. Structural changes, mutations, PTMs, and decreased synthesis of apolipoprotein A-I have been associated with multiple disease entities ^31^. In the context of liver disease, decreased apolipoprotein A-I levels have been observed in patients with cirrhosis ^32–36^. Although apolipoprotein A-I is a component of an established screening tool for liver fibrosis called the “Fibrotest,” scarce studies have assessed the ability of apolipoprotein A-I levels to predict decompensation events or death ^34–36^. In the most recent of these, Gurbuz et al used bottom-up proteomics to analyze the serum proteome of hepatocellular proteins in patients with cirrhosis versus healthy controls. Of 903 proteins identified, 81 proteins of hepatocellular origin were differentially expressed between groups: 65 downregulated and 16 upregulated in subjects with cirrhosis. Decreased serum levels of apolipoprotein A-I were observed in cirrhosis patients and correlated with a transition from stable to unstable decompensated cirrhosis ^36^. In contrast, we observed most apolipoprotein A-I proteoforms were upregulated in later stage disease. Modifications of apolipoprotein A-I proteoforms included truncation, oxidation of two methionine residues, and alpha-amino acetylation. Nearly all the observed proteoforms in later-stage disease (Figure S12) contained more amino acids compared to proteoforms seen in Stage I disease. This could be a result of secretion of apolipoprotein A-I’s prepropeptide and/or inappropriate cleavage of the prepropeptide in subjects with more severe disease ^31, 37^, ultimately leading to dysfunction of apolipoprotein A-I proteins. The hydroxylation of apolipoprotein A-I may exacerbate this as oxidation of HDL can negatively affect cholesterol efflux, anti-apoptotic effects, anti-peroxidation, and other anti-inflammatory actions ^38^. More specifically, oxidation of methionine residues within the apolipoprotein A-I protein can interfere with apolipoprotein A-I’s interaction with lecithin cholesterol acyltransferase (LCAT) impairing reverse cholesterol transport ^39^.

Haptoglobin, an acute-phase protein that scavenges cell-free hemoglobin, has also been studied as a biomarker of liver disease severity. Like apolipoprotein A-I, decreased haptoglobin levels are correlated with higher degrees of liver fibrosis ^40, 41^. There is also evidence to suggest glycoforms of haptoglobin may be useful in differentiating various stages and etiologies of liver disease ^41–47^. However, little is known about the specific effect of cirrhosis progression on haptoglobin function, and there are few to no studies that have investigated haptoglobin and its modifications as a predictor of future decompensation events. We observed DEPs of haptoglobin were relatively upregulated in Stage I disease (Figures 3, 4, S11). All haptoglobin DEPs were identified as a truncation, but review of the amino acid sequences would suggest DEPs derive from mature alpha-1 or alpha-2 haptoglobin chains (www.uniprot.org,^48^). This is additionally supported by the presence of two point mutations at residues 71 and 72, which are known mutations of two alleles of the alpha-1 chain, *alpha-1F* and *alpha-1S* ^48^. Interestingly, the alpha-amino acetylation PTM observed in our study was present only on the alpha-1 chain and has not been previously studied in the context of cirrhosis. Alpha-amino acetylation can impact protein half-life, protein folding, protein-protein interactions, and membrane targeting, but it is unknown what effect this specific PTM has on haptoglobin. As such, future investigations should elucidate the importance of this PTM in the setting of cirrhosis as it may impact the production of and function of mature haptoglobin proteins in patients with more severe disease.

Cirrhosis associated coagulopathy has been well described, and subjects with cirrhosis are predisposed to both bleeding and thrombotic events ^49^. Patients with stable liver disease generally have normal levels of fibrinogen, but as disease progresses fibrinogen levels steadily decrease ^50, 51^. Dysfibrinogenemia also plays an important role in coagulopathy as modifications to fibrinogen subunits can impact both structure and function ^52, 53^. In our cohort, fibrinogen DEPs originated most commonly from the alpha chain and were of varying lengths (S10-S12). Given the full-length fibrinogen alpha chain is ∼610 amino acids after translation and intracellular processing ^54^, it’s likely these alpha chain DEPs are byproducts of fibrin breakdown based upon their much shorter amino acid sequences (Figure S9). The discrepancies in length of detected DEPs across clusters may be a consequence of complex alterations in fibrin degradation and of the proteinases important for fibrin breakdown (*i.e.,* thrombin, plasmin, neutrophil elastase, and proteinase 3) in various stages of disease. This is consistent with studies that have shown D-dimer, a fibrin degradation product, is elevated in patients with cirrhosis and is associated with increased mortality in patients with later stage disease ^55–59^. Further characterization of these specific DEPs can provide a starting point for mechanistic investigations of fibrinolysis in patients with cirrhosis and may present an opportunity for therapeutic intervention, whether that be on the fibrinogen molecule itself or on the proteinases responsible for its degradation.

There are key limitations to our study. First, we are limited by the sample size of patients as this was an early pilot study to identify biomarker candidates. Considering this, a benefit of our proteoform-based analysis is the degree of resolution achieved, which can identify significant differences even in a small cohort study. Still, targeted validation studies in a larger cohort are needed to confirm and expand upon our findings. Second, our cohort consisted mostly of patients with alcohol or MASH-related cirrhosis, as they are the two most common etiologies of cirrhosis in the United States. Future studies will need to include additional patients with other etiologies of disease to ensure generalizability across cirrhosis populations. Finally, it is possible some proteoforms identified in our discovery TDP analysis are a result of electrospray ionization or sample handling during preparation. Specifically, oxidation is a known byproduct of such techniques, and we can assume that at least a few of the proteoforms detected may be a result of sample processing. However, the fact we observed significant differences in the expression of a number of oxidized proteoforms across samples that underwent identical processing makes this less likely and suggests biological processes are driving these differences.

In summary, quantitative TDP of plasma proteins from patients with cirrhosis revealed unique plasma proteoform profiles associated with stages of disease in this pilot study. To our knowledge, this is the first report of such proteoform analysis in patients with cirrhosis. Although we focused on the most abundant DEPs (apolipoprotein A-I, haptoglobin, and fibrinogen alpha chain), there were other PFRs identified in this study that may be candidate biomarkers for disease progression, both individually and in combination with other PFRs. Moving forward, we will perform targeted studies on larger cohorts to validate and further characterize the identified PFRs. The TDP methodology employed here could greatly increase the accuracy of blood-based analyses due to the identification of DEPs that are highly sensitive and specific probes of the underlying biology over time. In a clinical setting, understanding the change in PFRs from one stage of cirrhosis to the next would enable prompt recognition of patients transitioning from compensated to later-stage disease, presenting an opportunity for targeted risk reduction or early intervention in such patients.

## Abbreviations

BMI: (body mass index),
WBC: (white blood cell count),
HGB: (hemoglobin),
PLT: (platelets),
HCT: (hematocrit),
PT: (prothrombin time),
INR: (international normalized ratio),
AST: (aspartate aminotransferase),
ALT: (alanine aminotransferase),
ALP: (alkaline phosphatase),
TBILI: (total bilirubin),
CR: (serum creatinine),
Na: (Sodium),
GFR: (glomerular filtration rate),
MELD-Na: (Model for End Stage Liver Disease with Sodium),
CAP: (controlled attenuation parameter)
pHTN: (portal hypertension),
AIH: (autoimmune hepatitis),
PBC: (primary biliary cholangitis),
ETOH: (alcohol-associated liver disease),
HBV: (hepatitis B virus),
HCV: (hepatitis C virus),
MASH: (metabolic dysfunction associated steatohepatitis),
MASLD: (metabolic dysfunction steatotic liver disease),
pHTN: (portal hypertension),
HVPG: (Hepatic vein pressure gradients),
HE: (hepatic encephalopathy),
Top-down Proteomics: (TDP),
Differentially expressed proteoform: (DEP),
PTM: (post-translational modifications),
MOD: (Modification),
aa: (amino acids),
LC-MS/MS: (Liquid chromatography with tandem mass spectrometry),
PEPPI-MS: (Passively Eluting Proteins from Polyacrylamide gels as Intact species for MS),
GO: (gene ontology),
BP: (biological processes),
Cluster: (C),
HDL: (high-density lipoprotein),
LCAT: (lecithin cholesterol acyltransferase),
BCA: (bicinchoninic acid),
IRM: (ion routing multipole),
PAM: (Partitioning Around Medoids),
FDR: (false discovery rate).

## Acknowledgments

This study was supported by the National Institute of General Medical Sciences of the National Institutes of Health under P41GM108569 (NLK) and Transplant Innovation Endowment Grant (DL). JMS is supported by NIH Grant T32DK077662 and JMS and PP received support from the Steven J. Stryker, MD, Gastrointestinal Surgery Research and Education Endowment.

## Data Availability

Raw files and tdReport files can be found in Massive (Accession MSV000094311. The search results in the tdReport format can be viewed by using TDViewer freely available at http://topdownviewer.northwestern.edu. The search results were further analyzed, and figures were generated with R-custom scripts that are available on request.

## Supplemental data

Supplementary Figures were placed at the end of this document.

Supplementary Tables can be found here: https://doi.org/10.6084/m9.figshare.26062579

## Author Contributions

**Eleonora Forte**: Conceptualization, Investigation, Validation, Formal analysis, Writing-Original Draft, Visualization, Supervision, Project administration; **Jes M. Sanders:** Writing - Original Draft, Resources; **Indira Pla:** Software, Validation, Formal analysis, Data Curation, Writing-Original Draft, Visualization; **Vijaya Lakshmi Kanchustambham** and **Che-Fan Huang**: Methodology, Investigation, Validation, Writing - Review & Editing; **Michael A. R. Hollas**: Software, Validation, Formal analysis, Data Curation, Writing-Original Draft, Visualization; **Aniel Sanchez**: Investigation, Data Curation, Writing - Review & Editing; **Katrina N. Peterson:** Software, Validation, Formal analysis, Data Curation, Writing - Review & Editing; **Rafael D. Melani:** Conceptualization, Methodology, Writing – Review & Editing; **Alexander Huang:** Formal analysis, Writing – Review & Editing; Praneet Polineni, Julianna M. Doll, Zachary Dietch: Resources, Data Curation, Writing - Review & Editing, **Neil L. Kelleher:** Conceptualization, Writing – Review & Editing, Funding acquisition; Daniela P. Ladner: Conceptualization, Writing – Review & Editing, Funding acquisition.

## Supplementary Figures

**Figure S1.**
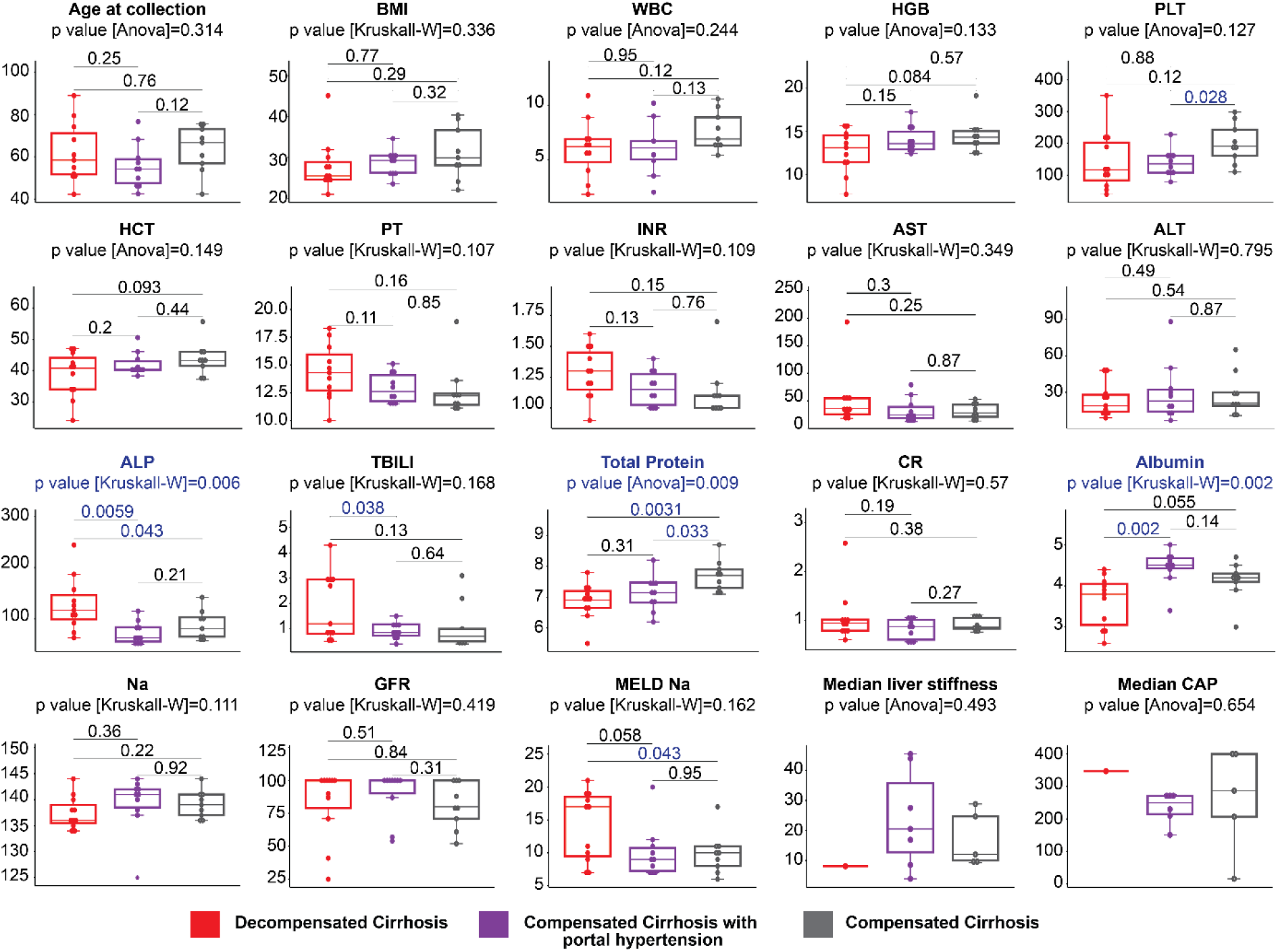
Clinical parameters of patients enrolled in this study. Clinical parameters shown in blue are statistically significant different among groups. Significance (p value <0.05) was calculated by Anova or Kruskall-Wallis (Kruskall-W) tests, depending on the data distribution determined by Shapiro test. Abbreviations: BMI (body mass index), WBC (white blood cell count), HGB (hemoglobin), PLT (platelets), HCT (hematocrit), PT (prothrombin time), INR (international normalized ratio), AST (aspartate aminotransferase), ALT (alanine aminotransferase), ALP (alkaline phosphatase), TBILI (total bilirubin), CR (serum creatinine), Na (Sodium), GFR (glomerular filtration rate), MELD-Na (Model for End Stage Liver Disease Sodium), CAP (controlled attenuation parameter).

**Figure S2.**
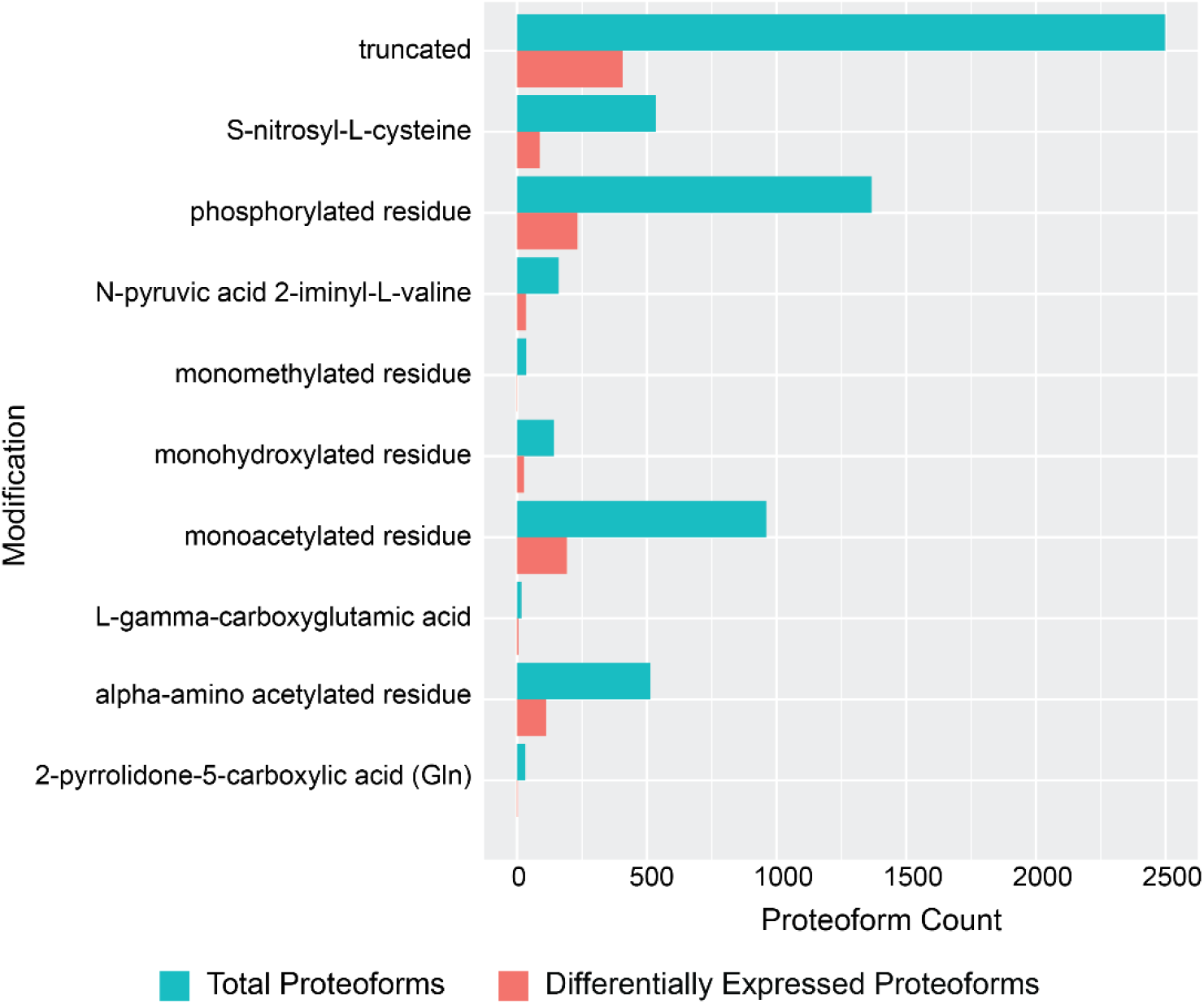
TCommon proteoform modifications detected in the TDP analysis. The number of truncations and top-10 most common post-translational modifications identified in the total and differentially expressed proteoforms (DEPs) captured in the discovery LC-MS/MS TDP analysis.

**Figure S3.**
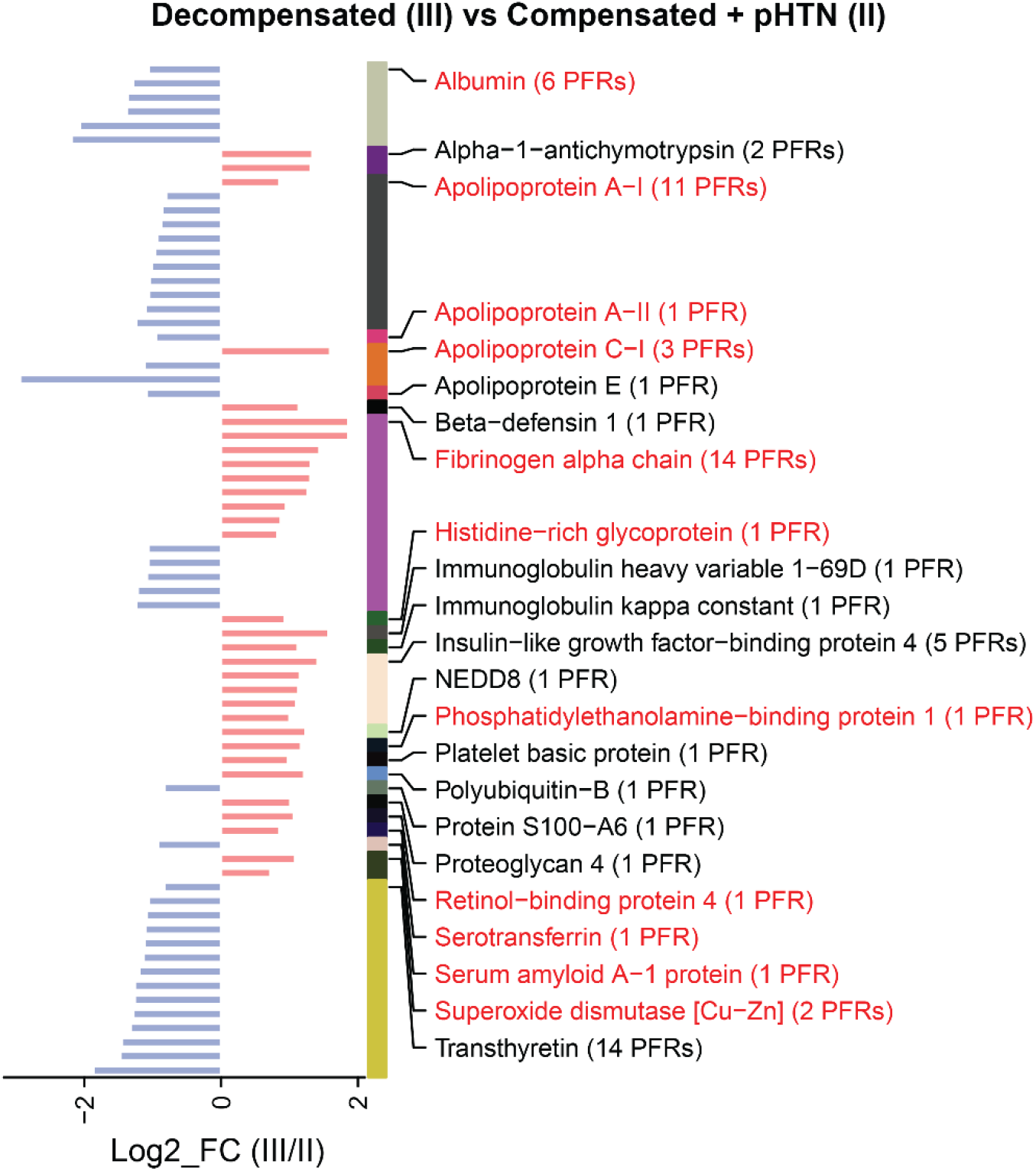
Differentially expressed proteoforms (DEPs) from decompensated (III) vs compensated with portal hypertension (+ pHTN) (II) patients. DEPs are grouped by proteins, and then ordered by fold change. The protein and number of DEPs derived from that protein are shown to figure right. For example, fibrinogen alpha chain had 14 DEPs identified. Proteins highlighted in red are enriched in liver at a transcriptional level. Abbreviations: PFR (proteoform).

**Figure S4..**
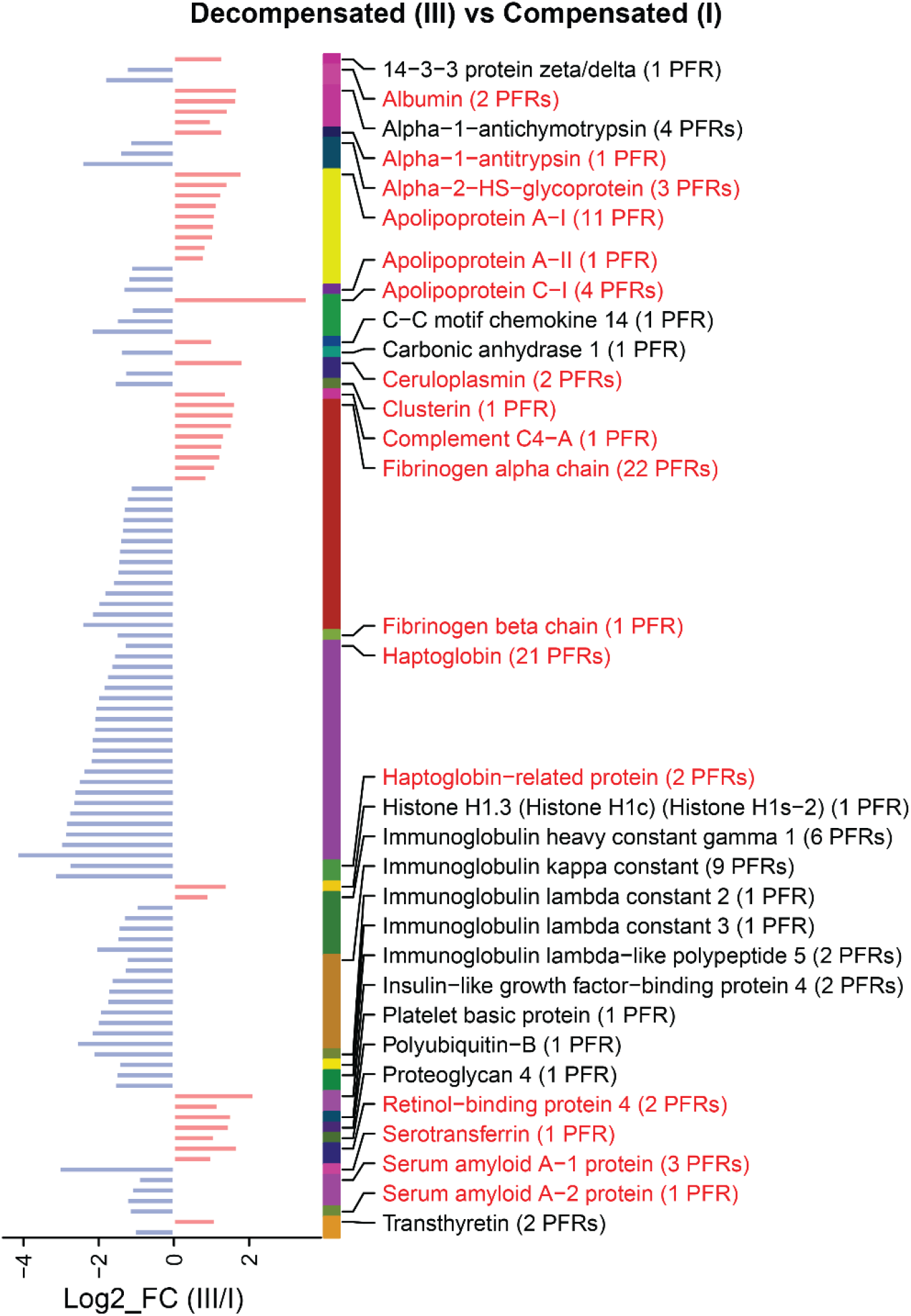
Differentially expressed proteoforms (DEPs) from decompensated (III) vs compensated (I) patients. DEPs are grouped by proteins, and then ordered by fold change. The protein and number of DEPs derived from that protein are shown to figure right. For example, haptoglobin related protein had 2 DEPs identified. Proteins highlighted in red are enriched in liver at a transcriptional level. Abbreviations: PFR (proteoform).

**Figure S5.**
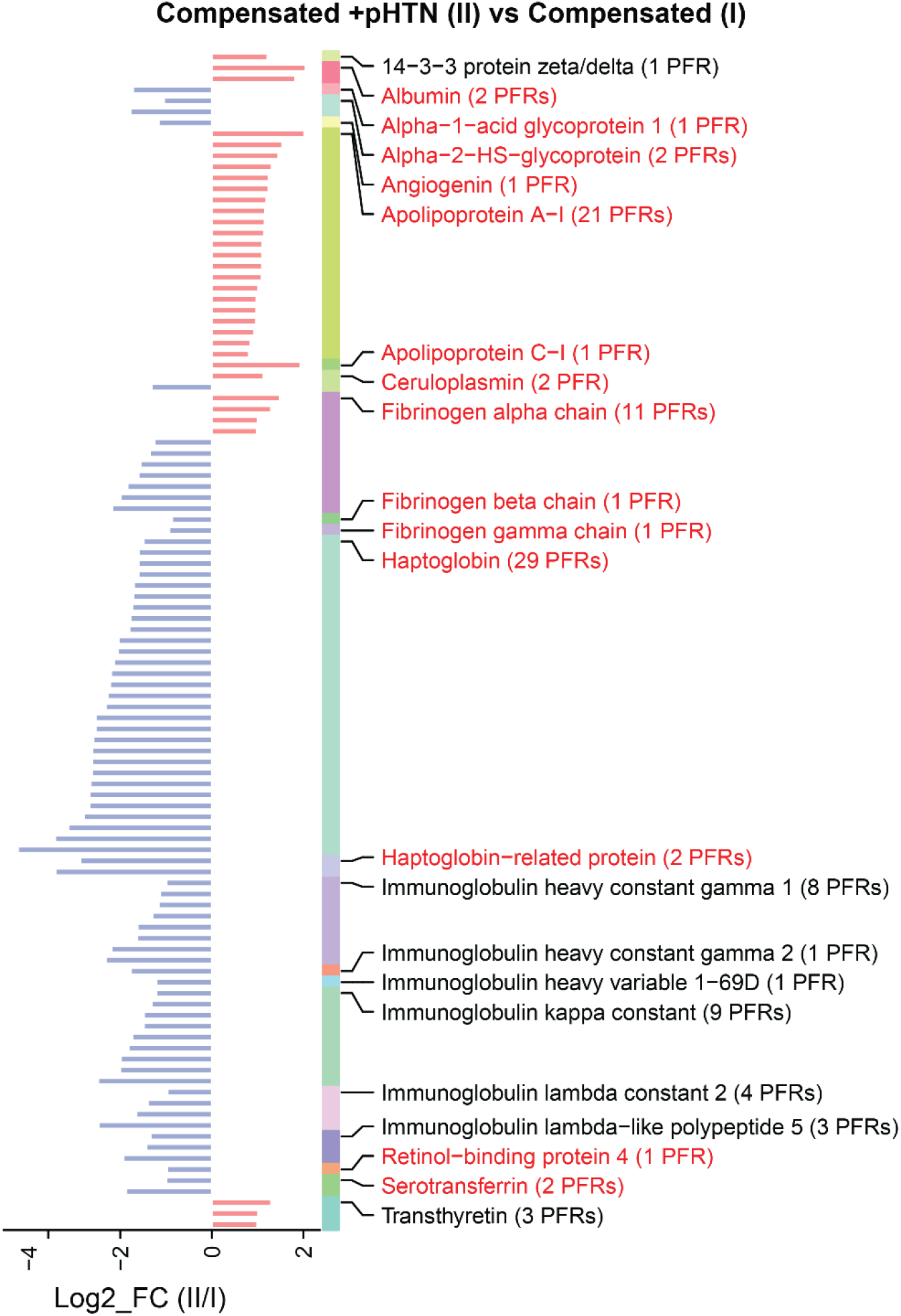
Differentially expressed proteoforms (DEPs) from compensated with portal hypertension (+ pHTN) (II) vs compensated (I) patients. DEPs are grouped by proteins, and then ordered by fold change. The protein and number of DEPs derived from that protein are shown to figure right. For example, haptoglobin related protein had 2 DEPs identified. Proteins highlighted in red are enriched in liver at a transcriptional level. Abbreviations: PFR (proteoform).

**Figure S6.**
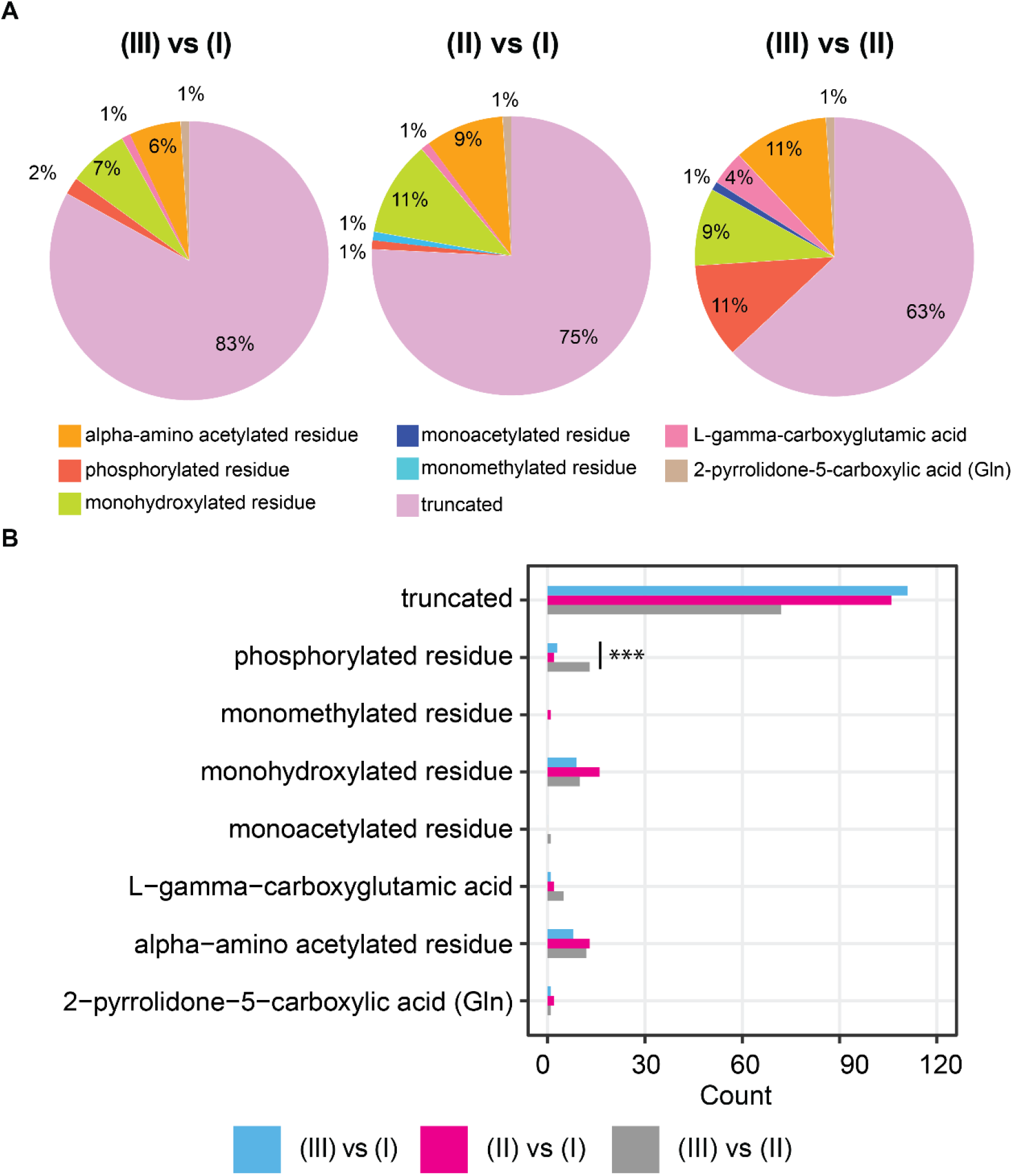
Proteoform modifications in pairwise comparisons of cirrhosis stages. **A)** Percentages and **B)** absolute number of differentially expressed proteoform (DEP) modifications identified in pairwise comparisons of Stage III vs I, II vs I, and III vs II. For example, 11% of proteoforms differentially expressed between Stages III and II were phosphorylated residues (red portion of pie chart). Statistical significance was calculated with the Fisher exact test (adj. *p*-values: ***<0.001, **<0.01, *<0.05).

**Figure S7.**
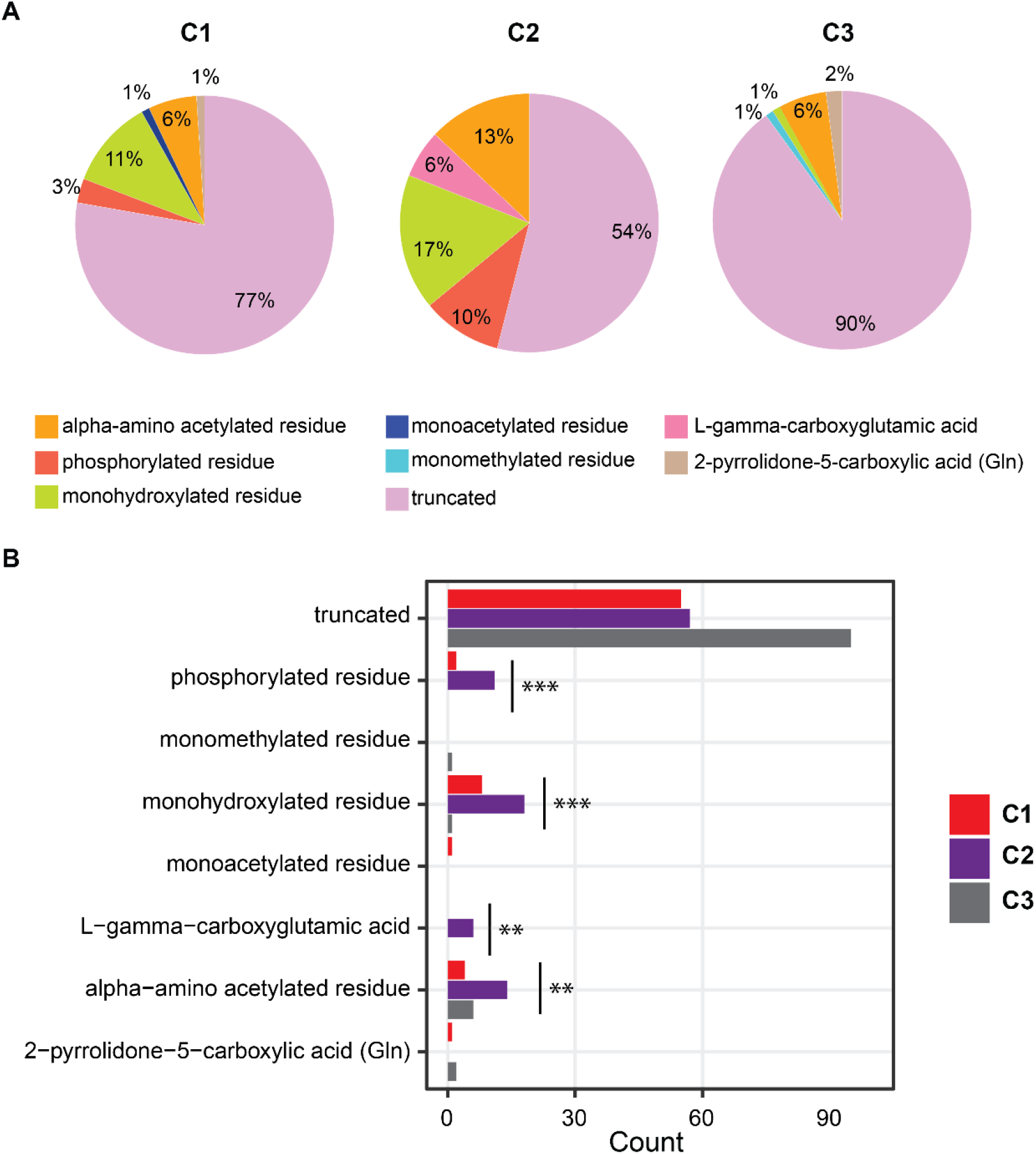
Proteoform modifications compared across clusters. **A)** Percentages and **B)** Absolute number of differentially expressed proteoform (DEP) modifications identified in clusters 1 (C1), 2 (C2), and C3. For example, 90% of modifications to DEPs that clustered into C3 were identified as truncations (light purple portion of the pie chart). Statistical significance was calculated with the Fisher exact test (adj. *p*-values: ***<0.001, **<0.01, *<0.05).

**Figure S8.**
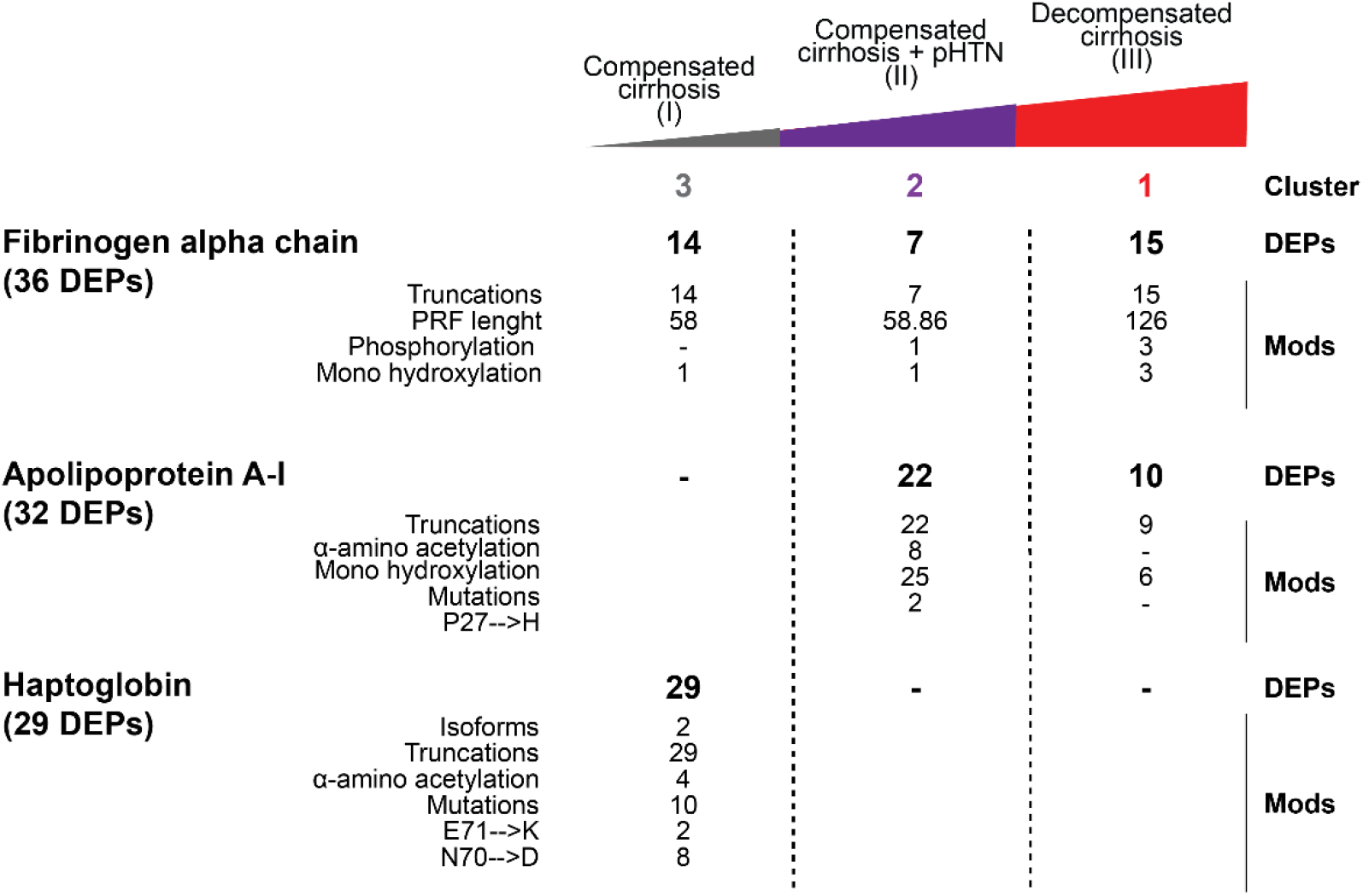
Most common, liver-enriched differentially expressed proteoforms (DEPs) identified in TDP analysis. DEPs relative to Fibrinogen alpha chain, Apolipoprotein A-I, and Haptoglobin are reported together with their modifications in the 3 clusters established in Figure 2. Proteins are shown with the number of DEPs in parentheses. Abbreviations: DEP (Differentially expressed proteoform), Mod (Modification).

**Figure S9.**
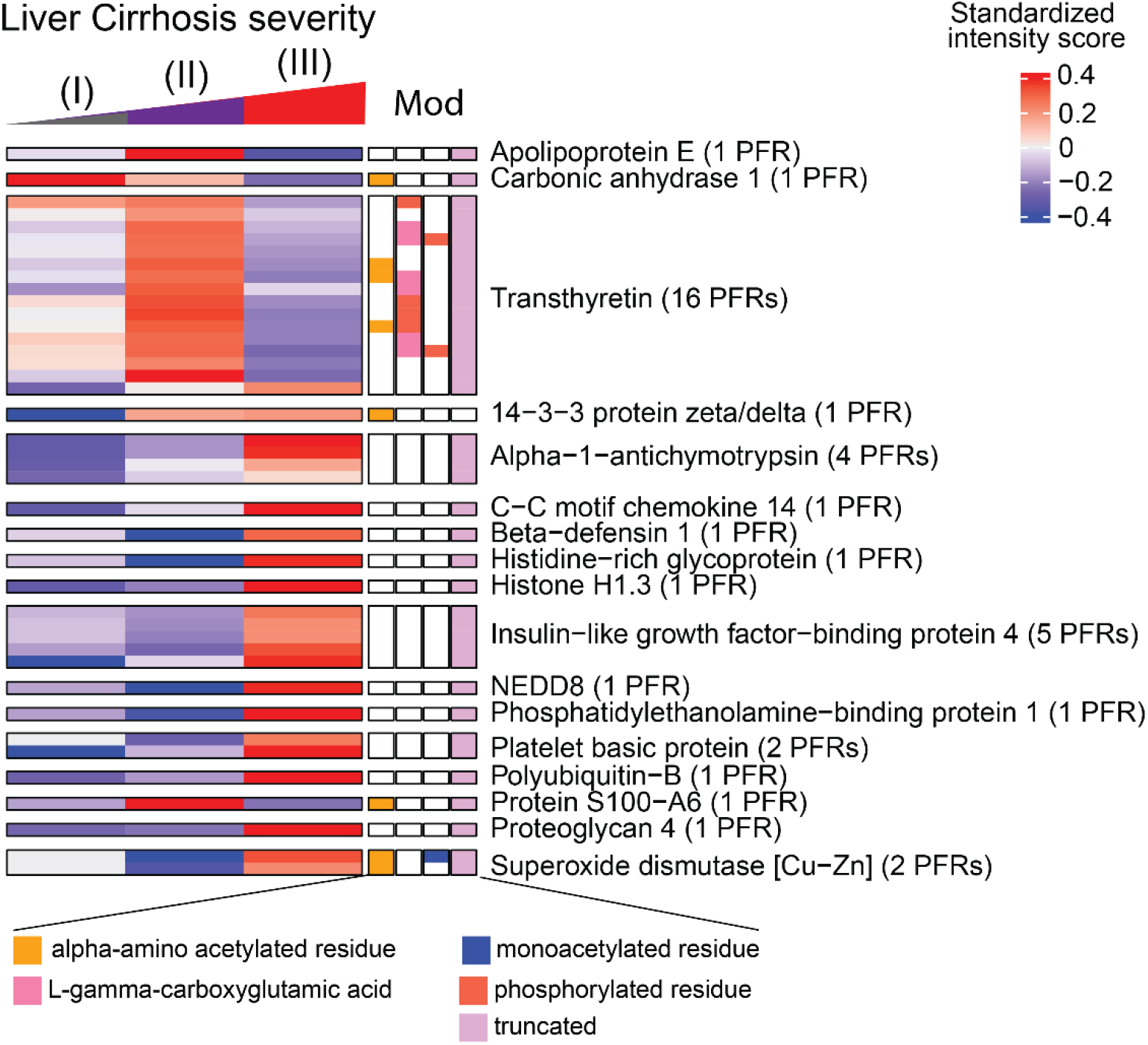
Heatmaps of quantified proteoforms of proteins not enriched in the liver at transcriptional level. Associated proteoform modifications are shown and defined by the figure legend. The proteins and number of identified proteoforms derived from each protein are shown to figure right. Abbreviations: PFR (proteoform), Mod (modification).

**Figure S10.**
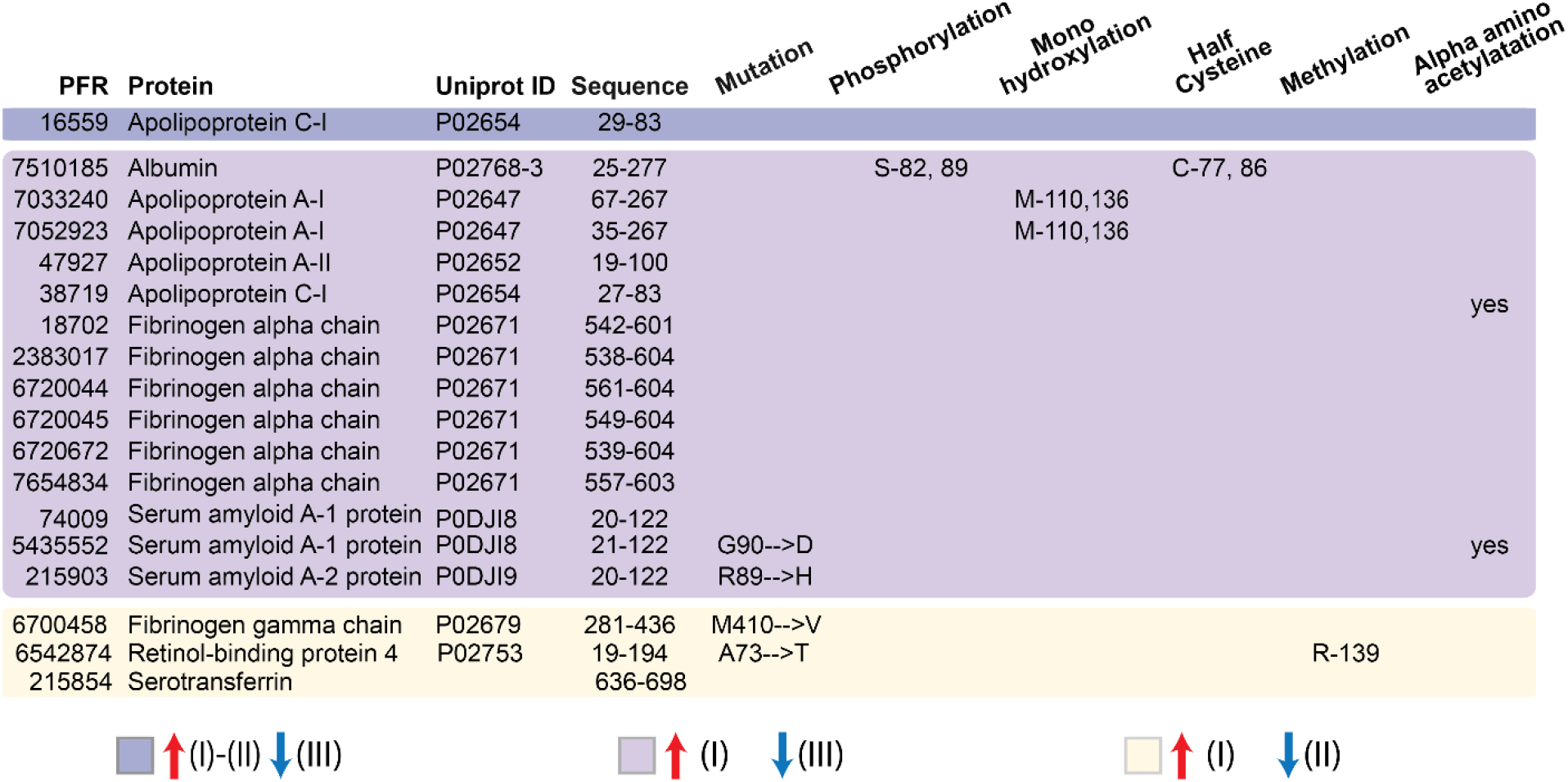
Individual proteoforms upregulated in early-stage cirrhosis. Differentially expressed proteoforms (DEPs) significantly upregulated in stages (I) and (II) and downregulated in stage (III) (dark purple), upregulated in stage (I) and downregulated in stage (III) (light purple), upregulated in stage (I) and downregulated in stage (II) (yellow). Each proteoform is shown with its unique proteoform number, protein name,Uniprot ID, amino acid sequence relative to the Uniprot ID, any relevant mutations, and presence or absence of post-translational modifications. Colors are based on the proteoform signatures created in Figure 4. Abbreviations: PFR (proteoform). *Note that half Cysteine could be artifact during the identification process.

**Figure S11.**
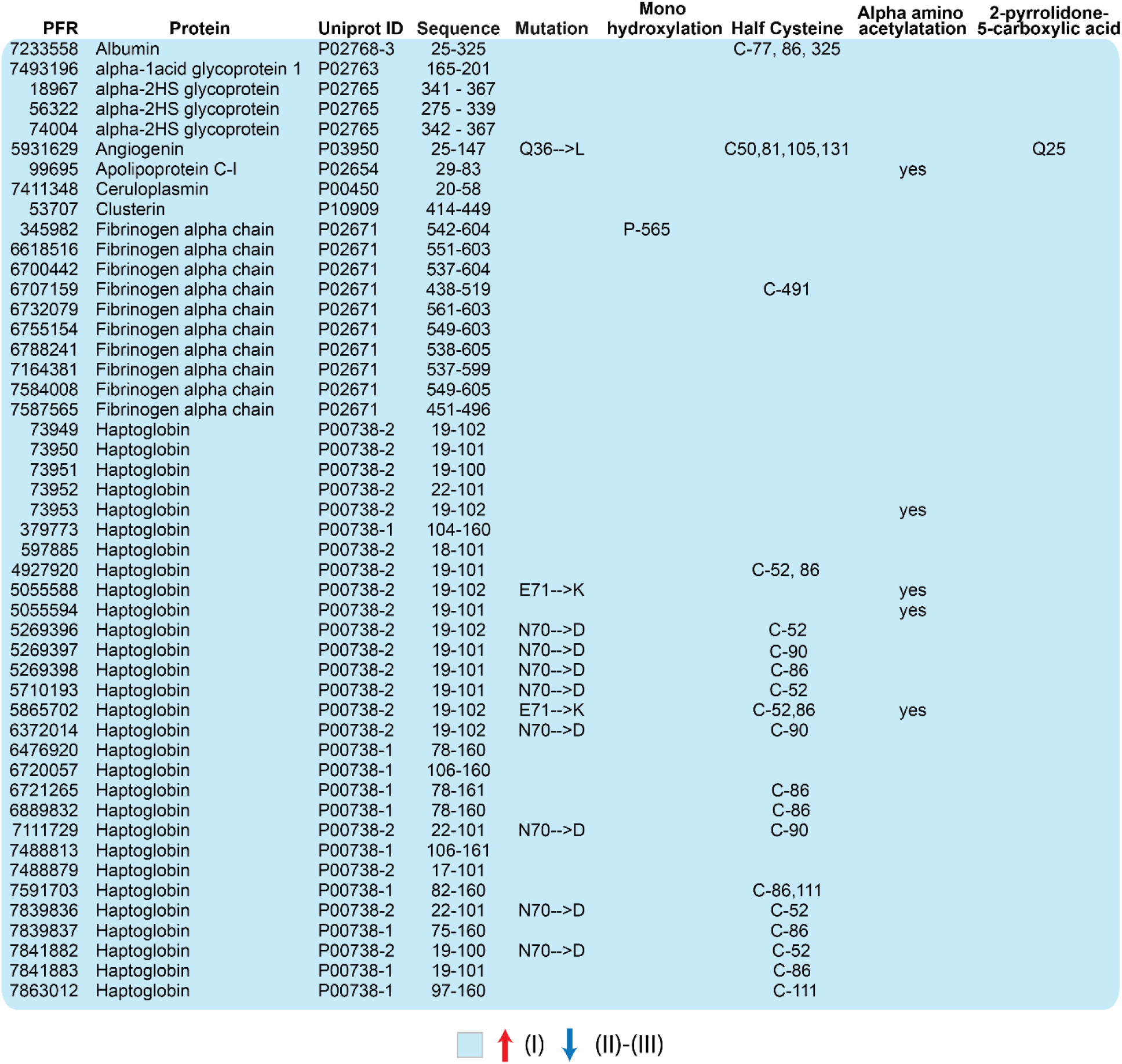
Individual proteoforms upregulated in Stage I disease. Differentially expressed proteoforms (DEPs) significantly upregulated in stage (I) and downregulated in stages (II) and (III) (blue). Each proteoform is shown with its unique proteoform number, protein name,Uniprot ID, amino acid sequence relative to the Uniprot ID, any relevant mutations, and presence or absence of post-translational modifications. Colors are based on the proteoform signatures created in Figure 4. Abbreviations: PFR (proteoform). *Note that half Cysteine could be artifact during the identification process.

**Figure S12.**
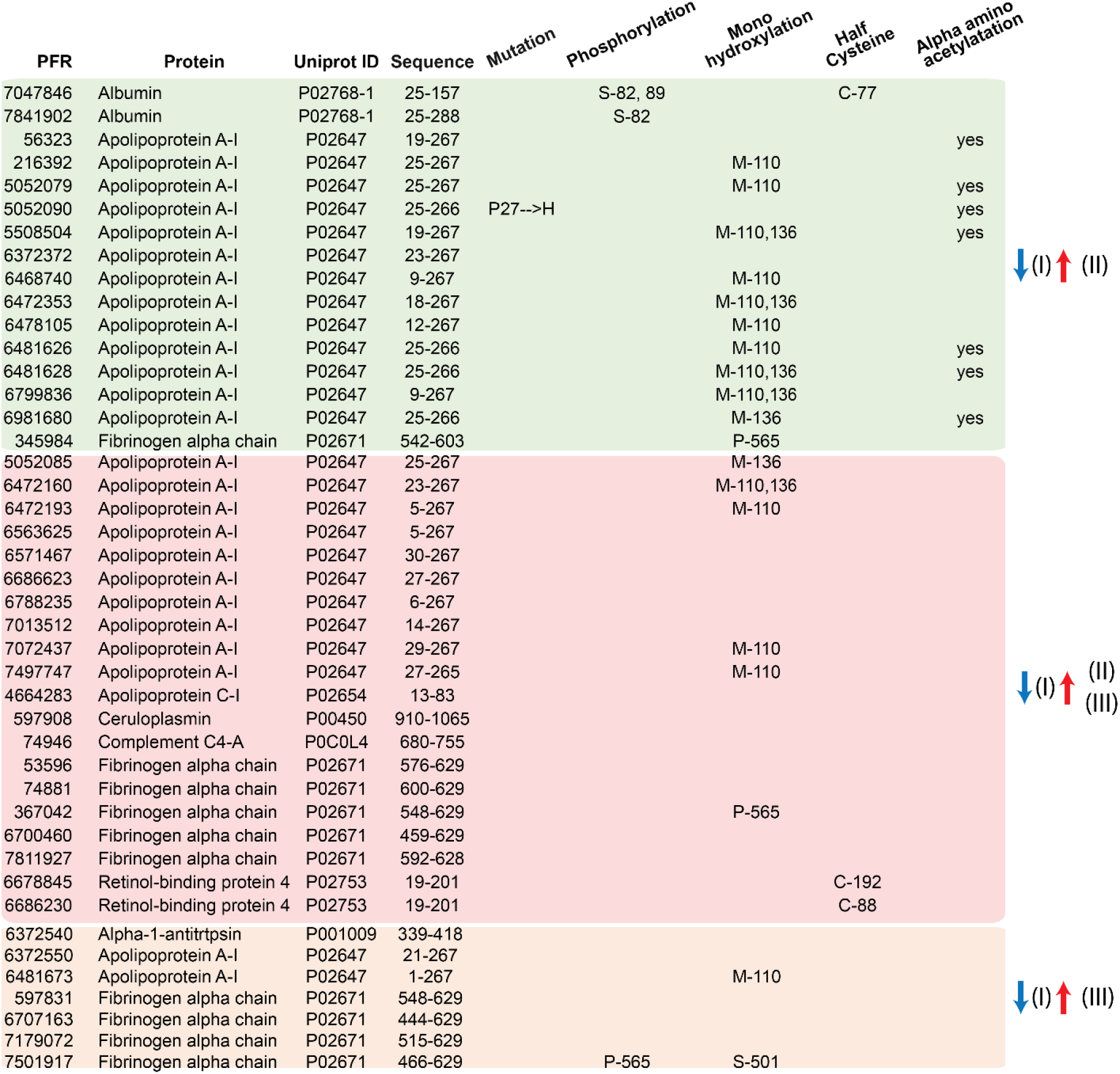
Individual proteoforms upregulated in late-stage cirrhosis. Differentially expressed proteoforms (DEPs) significantly downregulated in stage (I) and upregulated in stage (II) (green), downregulated in stage (I) and upregulated in stages (II) and (III) (red), downregulated in stage (I) and upregulated in stage (III) (orange). Each proteoform is shown with its unique proteoform number, protein name and Uniprot ID, amino acid sequence relative to the Uniprot ID, and presence or absence of post-translational modifications. Colors are based on the proteoform signatures created in Figure 4. Abbreviations: PFR (proteoform). *Note that half Cysteine could be artifact

